# Evidence for an Active Role of Inferior Frontal Cortex in Conscious Experience

**DOI:** 10.1101/2020.05.28.114645

**Authors:** Veith Weilnhammer, Merve Fritsch, Meera Chikermane, Anna-Lena Eckert, Katharina Kanthak, Heiner Stuke, Jakob Kaminski, Philipp Sterzer

## Abstract

2

In the search for the neural correlates of consciousness, it has remained controversial whether prefrontal cortex determines what is consciously experienced or, alternatively, serves only complementary functions such as introspection or action.

Here, we provide converging evidence from computational modeling and two functional magnetic resonance imaging experiments for a key role of inferior frontal cortex in detecting perceptual conflicts that emerge from ambiguous sensory information. Crucially, the detection of perceptual conflicts by prefrontal cortex turned out to be critical in the process of transforming ambiguous sensory information into unambiguous conscious experiences: In a third experiment, disruption of neural activity in inferior frontal cortex through transcranial magnetic stimulation slowed down the updating of conscious experience that occurs in response to perceptual conflicts.

These findings show that inferior frontal cortex actively contributes to the resolution of perceptual ambiguities. Prefrontal cortex is thus causally involved in determining the contents of conscious experience.

**One-sentence Summary:** Inferior frontal cortex detects and resolves perceptual conflict during bistable perception.

## 4 Introduction

The neural basis of conscious experience is one of today’s greatest mysteries^1^. Its unravelling will have important implications for how we approach patients who remain unresponsive after brain damage or who suffer from hallucinatory distortions of perception^2^. Likewise, such progress may expand our ability to detect the presence of conscious experience in organic and artificial life beyond the human mind^3^. Yet, solutions to these problems remain elusive, since they require identifying not only the *neuro-anatomic correlates* of consciousness, but also the *computational function* of specific brain regions for conscious experience^4,5^.

In the search for such a neuro-computational understanding of consciousness, the role of prefrontal brain areas is currently one of the most debated topics: While some theories posit that prefrontal cortex plays a critical role in perceptual consciousness^4^, others have allocated the neural correlates of conscious experience to more posterior brain regions^5^. The latter accounts link prefrontal activity to cognitive processes that occur in the wake of conscious experience^6^, such as evaluating or acting on the contents of perception^7^.

Bistable perception has been a key experimental approach in this debate for more than two decades^8^. In this phenomenon, stimuli that are compatible with two interpretations give rise to *perceptual conflict*^9^. Faced with this conflict, observers perceive periodic changes in conscious experience, oscillating between two mutually exclusive perceptual states^10^. Thereby, bistable perception provides a unique window into a fundamental neuro-computational aspect of consciousness: the transformation of ambiguous sensory information into unambiguous conscious experience^11,12^.

Functional neuroimaging studies in humans have identified the right inferior frontal cortex (IFC; a subregion of prefrontal cortex, Figure 1A) as a key region in bistable perception^8^. When compared to stimulus-driven changes in perception, spontaneous perceptual changes during bistability were consistently associated with increased activity in IFC^8^, suggesting that prefrontal cortex actively contributes to conscious experience^10,12–14^. Along this line of thought, IFC may resolve perceptual conflict by triggering perceptual changes through feedback signaling to sensory areas^10,12^ (Figure 1A).

**Figure 1.**
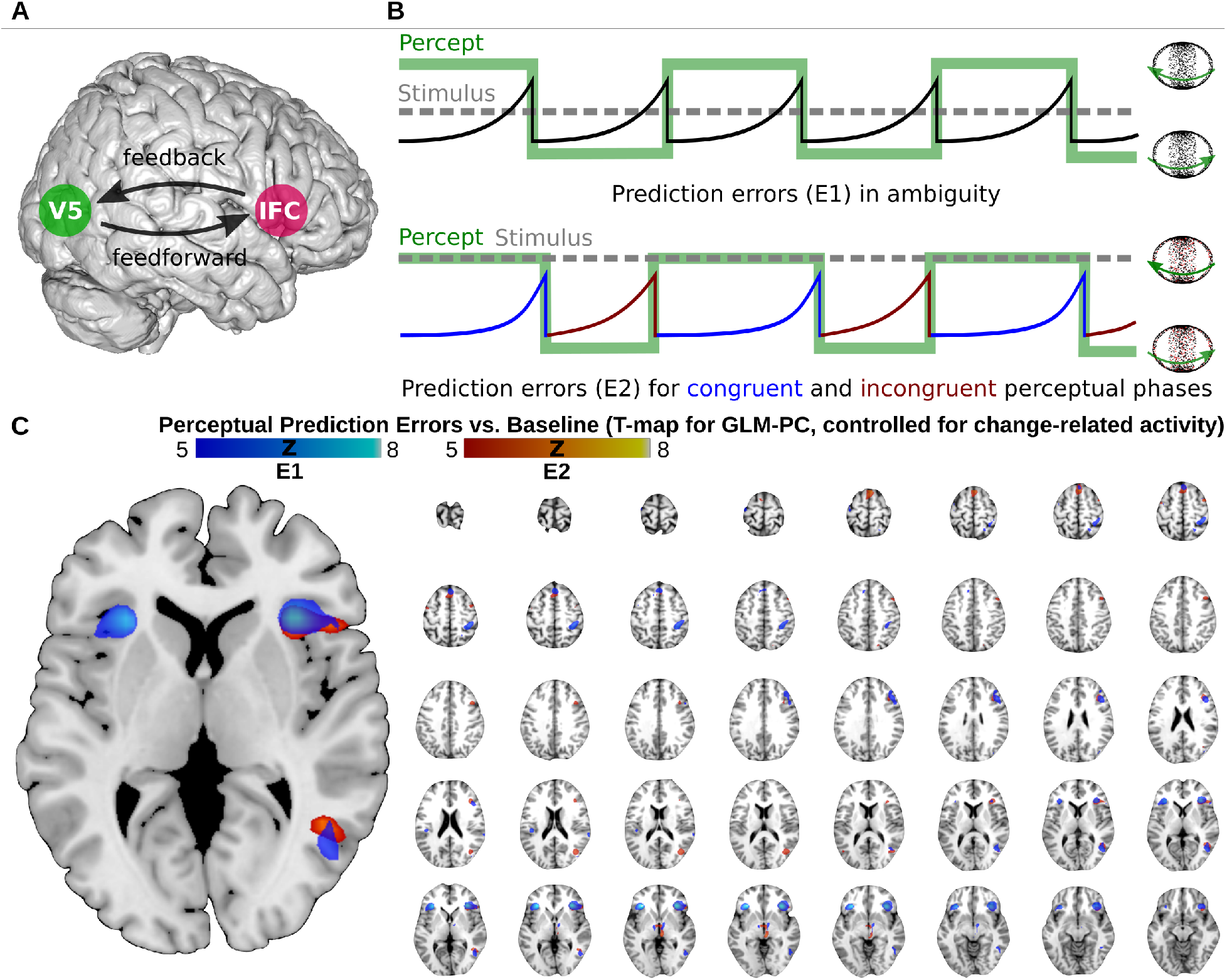
**A. IFC in bistable perception.** The role of IFC (inferior frontal cortex, schematic overlay in pink) in conscious experience is controversial: According to one view, feedback from IFC may modulate perceptual processing in visual cortex (motion-sensitive visual cortex V5/hMT+, highlighted in green). This may reflect an active contribution of prefrontal brain activity to conscious experience. The opposing view links neural activity in IFC to the quality, report or reportability of perceptual events. This suggests that conscious experience may be realized within visual cortex and may drive activity in IFC by means of feedforward processing. **B. Accumulating perceptual conflict and conscious experience.** Here, we depict conscious experience and associated changes in accumulating perceptual conflict for bistable perception induced by a random dot structure-from-motion stimulus (RDK). Perceived direction of rotation (green line) alternates between left- and rightward motion of the front-surface (icons on the right). In the absence of disambiguating sensory evidence (upper panel; grey dotted line), prediction errors (black solid line) accumulate while perception remains constant, until perceptual conflict is reduced by a change in conscious experience. When faced with additional stimulus information (lower panel), conscious experience fluctuates between *congruent* or *incongruent* perceptual states. If an observer adopts a percept that is *congruent* with the disambiguating stimulus information, prediction errors are reduced (blue line). Accordingly, conflict-driven changes in conscious experience toward the alternative stimulus interpretation become less likely (vice versa for incongruent perceptual states, red line). **C. Neural correlates of accumlating perceptual conflict.** Converging evidence from two experiments (E1: blue heatmap; E2: red heatmap; both displayed for T > 5) indicates that perceptual prediction errors correlate with neural activity in right IFC (anterior insula and inferior frontal gyrus) and V5/hMT+, while controlling for change-related BOLD responses. Additional activations were located in left insula, right posterior-medial frontal gyrus and right inferior parietal lobulus (all *p_FWE_* < 0.05, see **corresponding Supplementary Table S1**).

However, this *feedback-account* has been questioned by work that related IFC activity to the quality^15^, report^16^ and reportability^17,18^ of changes in conscious experience during bistable perception. Perceptual conflict may thus rather be resolved within visual cortex and elicit activity in IFC through a *feedforward* mechanism. Accordingly, IFC activity may not reflect the cause, but the consequence of changes in conscious experience.

To settle the ongoing controversy between these feedforward- and feedback-accounts of bistable perception will be a critical step in elucidating the role of prefrontal cortex in consciousness. Here, we conjectured that these seemingly contradictory views may be reconciled within one computational characterization of IFC that incorporates both feedforward and feedback processing.

Firstly, we hypothesized that IFC may signal perceptual conflicts that arise between the contents of conscious experience and the available sensory data through a feedforward mechanism. To test this hypothesis, we used functional magnetic resonance imaging (fMRI) in conjunction with computational modeling in a Bayesian framework. Secondly, we reasoned that the detection of perceptual conflict by IFC may in turn trigger changes in conscious experience via feedback signaling to sensory areas. This latter hypothesis was tested by disrupting IFC activity through neuronavigated transcranial magnetic stimulation (TMS).

## 5 Results

In a series of three experiments (E1-E3, Supplementary Figure S1), human observers reported changes in their perception of a rotating sphere (rightward-, leftward or unclear direction of rotation). In this *structure-from-motion* stimulus, random dots distributed on two intersecting rings induce illusory 3D motion (see Supplementary Video 1 and Supplementary Materials 9.1). Due to the perceptual conflict inherent in the ambiguous visual input, participants perceived spontaneous changes between left- and rightward rotation that occurred every 25.08 ± 2.57 sec (*phase duration*, i.e., the average time spent between two consecutive changes in conscious experience; Supplementary Figure S2A-B).

With this type of stimulus, unclear perceptual states^15^ are rare (0.6 ± 0.18%). Moreover, changes in perceived direction of rotation occur only when fore- and background of the illusory sphere overlap^19^ (Supplementary Figure S3A-B). These perceptual features ensured the temporal precision of our approach and allowed us to compute response times (average *RT* = 0.81 ± 0.05 sec) as a control variable for processes associated with behavioral reports^16,17^.

### 5.1 IFC detects accumulating perceptual conflict

Bayesian perceptual inference^20^ provides a plausible computational explanation for the effects of conflicting stimulus information on perception. In this framework, conscious experience is understood as an iterative process, constantly generating and testing *hypotheses* about the most likely cause of sensory data^21^. In bistable perception, the two alternating stimulus interpretations reflect mutually exclusive hypotheses that are both compatible with, but cannot fully account for the ambiguous sensory data. This results in perceptual conflict^8,9,12^. As a quantitative representation of such conflict, the residual evidence for the alternative to the currently dominant stimulus interpretation can be conceived of as a *perceptual prediction error*^11^. This unexplained error induces a progressive shift in the balance between the two perceptual hypotheses^14^. Over time, prediction errors therefore accumulate until the increasing perceptual conflict is reduced by a change in conscious experience from the dominant to the alternative stimulus interpretation (Figure 1B). A recent proof-of-concept study has linked this process to neural activity in prefrontal cortex^14^: During ambiguous visual stimulation, BOLD signals in IFC gradually increased while perception remained constant, peaking at the time of a conflict-induced change in conscious experience.

In experiment E1 (Supplementary Figure S1A), we sought to (1) confirm the previously suggested role of IFC in detecting perceptual conflict and to (2) test the hypothesis that such perceptual conflict originates from visual cortex.

To identify the neural representation of perceptual conflict, we acquired fMRI data during bistable perception and searched for correlations of BOLD activity with the dynamics of perceptual prediction errors. To this end, we inverted an established computational model of bistable perception^11,14^ (see Supplementary Materials 9.2) based on the individual participants’ behavioral reports on perceptual changes. This model estimates dynamic perceptual prediction errors to explain the sequence of conscious experiences during bistable perception. In model-based fMRI, we searched for the neural correlates of these prediction errors while controlling for BOLD activity associated with the timing and report of perceptual changes. In line with previous results^14^, we found that perceptual prediction errors correlated with BOLD activity in right-hemispheric IFC (anterior insula, inferior frontal gyrus^8^; Bonferroni-corrected for family-wise error *p_FWE_* < 0.05, Figure 1C, Supplementary Table 1).

Of note, previous studies have predominantly linked IFC to event-related processes associated specifically with perceptual changes^8^. We reasoned that this often-reported finding of change-related activity in IFC may actually correspond to the peak of accumulating prediction errors (see Figure 1B and Supplementary Figure S4). In our data, such change-related IFC activity was only detectable if the analysis did not control for prediction errors (Supplementary Figure S5). Indeed, a direct comparison based on posterior probability maps^22^ confirmed that BOLD activity in right-hemispheric IFC was better explained by prediction errors that gradually accumulated in each perceptual phase than by perceptual change events (Supplementary Figure S6).

This suggests that, during bistable perception, IFC activity reflects the gradual accumulation of perceptual conflict^14^, until it is temporarily reduced by a conflict-driven change in conscious experience (Figure 1B, see below for a replication of this finding in experiment E2).

Yet, as a supra-modal brain region, IFC is unlikely to represent visual information independently of sensory brain regions. We therefore hypothesized that information about perceptual conflict may be fed forward to IFC from the representation of perceptual content in visual cortex^23^. Indeed, perceptual prediction errors also correlated with BOLD activity in the motion-sensitive extrastriate visual area V5/hMT+^24^ (Figure 1C). Dynamic causal modeling^25^ confirmed that these signals of accumulating perceptual conflict were most likely to originate from V5/hMT+, reaching IFC via feedforward effective connectivity (Supplementary Figure S7).

In contrast to IFC, neural activity in V5/hMT+ also reflected the content of conscious experience, that is, whether participants experienced leftward or rightward rotation during bistable perception: Based on multi-voxel pattern analysis^26^ of BOLD signals in V5/hMT+, we were able to decode which of the two stimulus interpretations was, at a given point in time, *dominant* or *suppressed* (Supplementary Materials 9.3 and Supplementary Figure S8).

We therefore asked whether the neural correlates of perceptual conflict originated from the voxels that representated the currently suppressed stimulus interpretation in visual cortex. As predicted, the BOLD signal in V5/hMT+ voxels that showed enhanced activity during perception of *leftward* rotation correlated more strongly with perceptual prediction errors when participants experienced *rightward* rotation (*BF*_10_ = 2.24 × 10^3^, Figure 2A) and vice versa (*BF*_10_ = 2.57 × 10^3^, see below for a replication of this finding in E2). This intriguing dissociation between the representation of perceptual content and perceptual conflict occurred only in voxels with strong biases toward one of the two stimulus interpretations (Supplementary Figure S9A).

**Figure 2.**
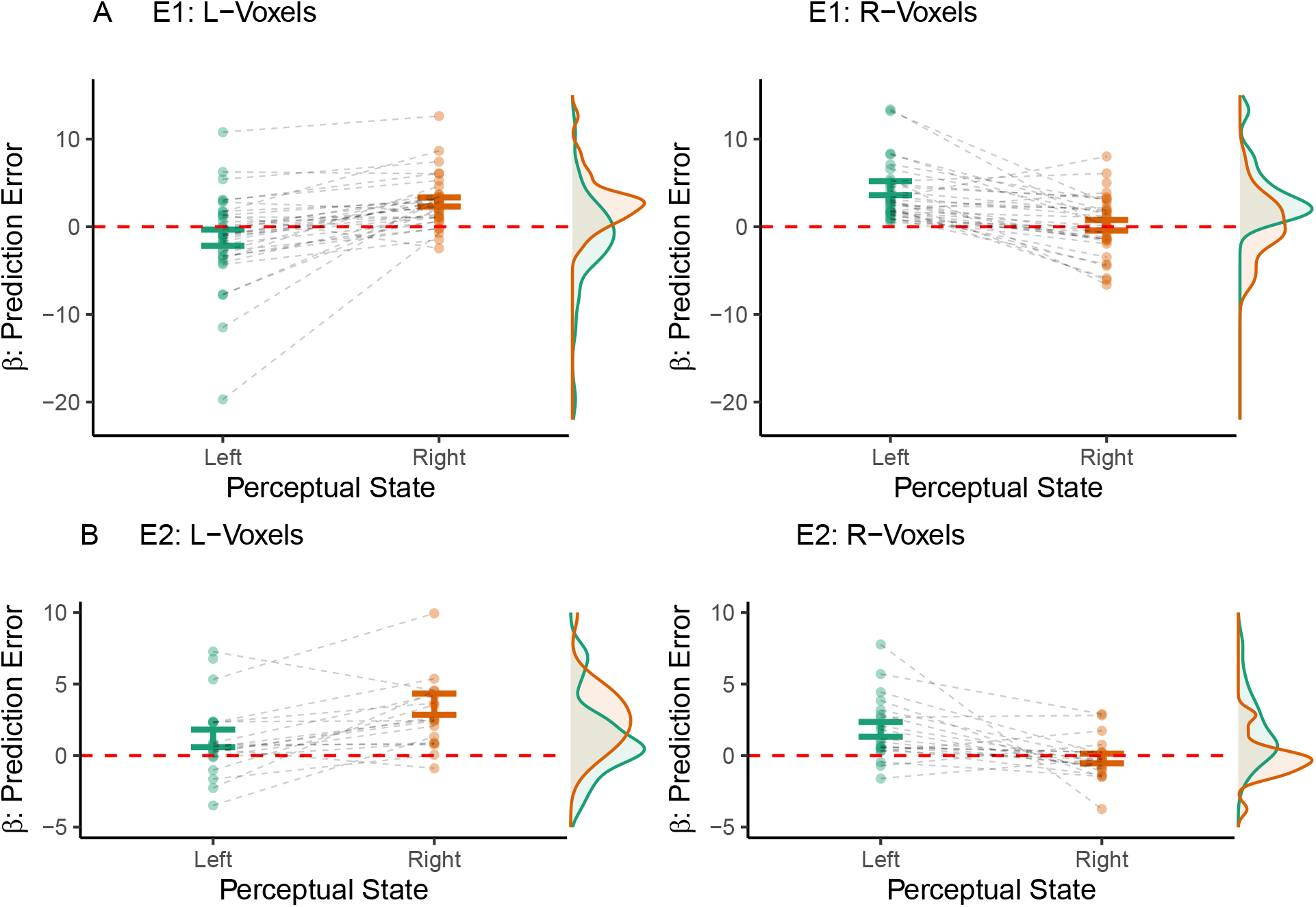
Neural correlates of perceptual conflict in V5/hMT+. **A. Experiment E1.** We delineated V5/hMT+ based on the effects of visual stimulation (i.e., independently of our computational model of bistable perception) and identified *biased* voxel populations that showed elevated neural activity during either leftward (L) or rightward (R) illusory rotation (T-value > 1; average number of voxels per population *N_pop_* = 33.97 ± 1.78). While controlling for effects related to perceptual changes, we found that BOLD activity in L-voxels (left panel) correlated more strongly with perceptual prediction errors when participants consciously perceived rightward rotation (paired t-test: T(32) = −5.26, p = 9.22 × 10^-6^, *BF*_10_ = 2.24 × 10^3^). Conversely, R-voxels (right panel) correlated more strongly with perceptual prediction errors when participants consciously perceived leftward rotation (T(32) = 5.32, p = 7.94 × 10^-6^, *BF*_10_ = 2.57 × 10^3^). **B. Experiment E2.** Experiment E2 (*N_pop_* = 32.92 ± 3.12) replicated these results: L-voxels (left panel) correlated more strongly with perceptual prediction errors during illusory rotation toward the right (paired t-test: T(19) = −4.07, p = 6.49 × 10^-4^, *BF*_10_ = 53.36). Inversely, R-voxels (right panel) correlated more strongly with perceptual prediction errors when the participants consciously perceived leftward rotation (T(19) = 3.11, p = 5.71 × 10^-3^, *BF*_10_ = 8.2).

### 5.2 IFC is sensitive to graded changes in perceptual conflict

The results of E1 indicate that IFC receives feedforward information about perceptual conflict, emanating from the representations of ambiguous stimuli in visual cortex. Yet, in everyday perception, fully ambiguous stimuli like those giving rise to bistable perception are rare. Rather, additional (i.e., *disambiguating*) stimulus information is usually available, albeit often incomplete^27^. In experiment E2, we sought to confirm the role of IFC in the signaling of perceptual conflict by measuring its responses to such disambiguating stimulus information.

To this end, we conducted an independent fMRI-experiment (Supplementary Figure S1B) based on the novel paradigm of *graded ambiguity*^24,28^. As in E1, participants reported changes in the perceived direction of rotation of a structure-from-motion stimulus. In contrast to E1, we introduced random changes in a disambiguating 3D signal attached to a fraction of the stimulus dots. The amount of disambiguating information was varied parametrically across six levels of *signal-to-ambiguity ratio*. As a consequence, conscious experience fluctuated to varying degrees between perceptual states that were *congruent* or *incongruent* with the disambiguating stimulus information (Figure 1B, lower panel). We assumed that, depending on the signal-to-ambiguity ratio, perceptual conflict should be greater during incongruent perceptual states, thus increasing the likelihood of conflict-driven changes toward the alternative stimulus interpretation.

As expected^28^, congruent perceptual states were indeed more frequent for increasing signal-to-ambiguity ratios (*BF*_10_ = 2.89 × 10^22^, Figure 3A). Both model simulation (Supplementary Figure S10) and computational modeling of the participants’ behavior (Figure 3B) confirmed that prediction errors were enhanced during incongruent as compared to congruent perceptual states (main effect of *Congruency*), with stronger effects at higher signal-to-ambiguity ratios (interaction between *Congruency* and *Signal-to-Ambiguity*).

**Figure 3.**
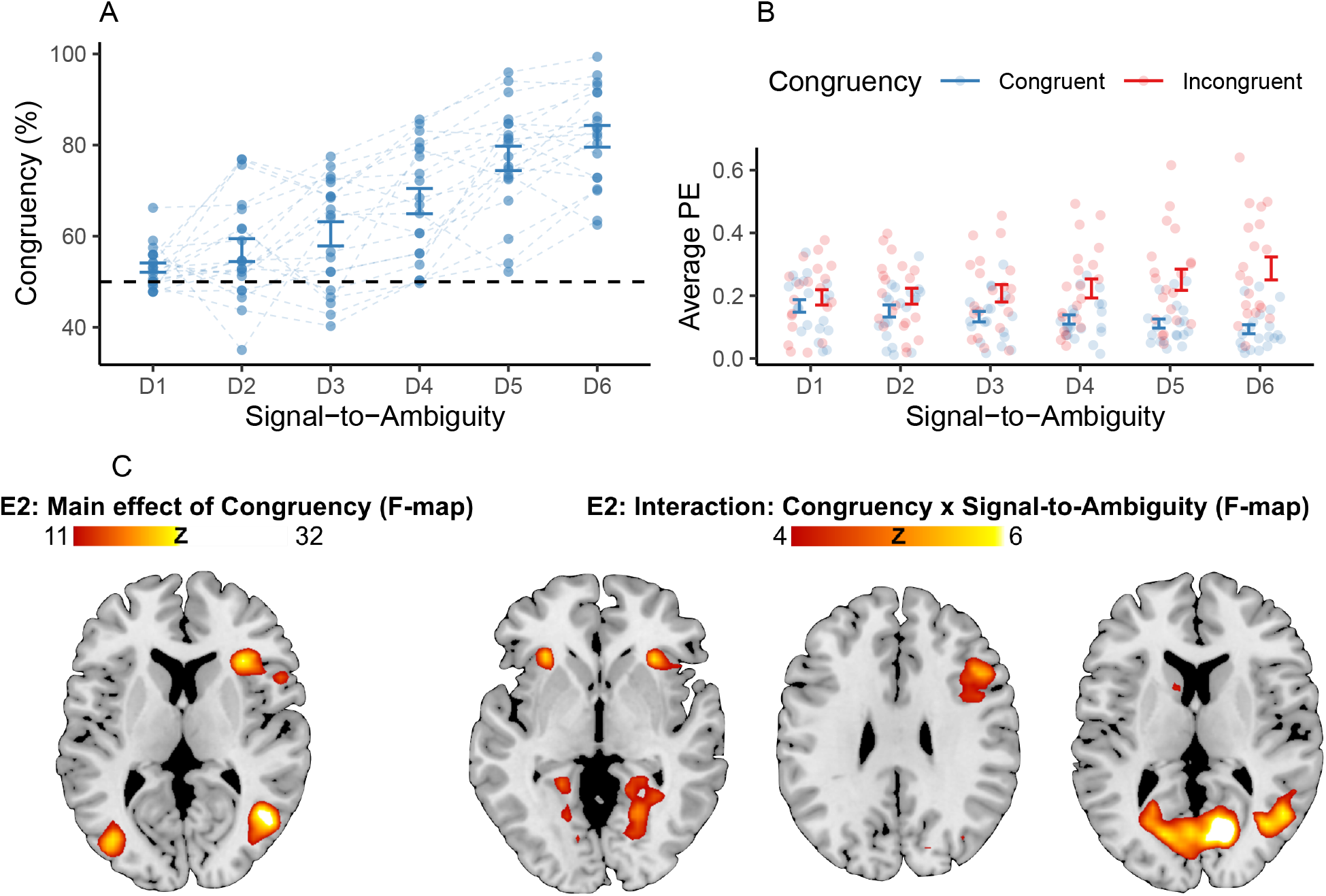
Perceptual conflict during graded ambiguity. **A. Congruent perceptual states.** Conscious experience was biased toward perceptual states that were congruent with the disambiguating stimulus information (T(19) = 8.45, p = 7.37 × 10^-8^, *BF*_10_ = 1.97 × 10^5^). For increasing signal-to-ambiguity ratios (levels D1 to D6), congruent perceptual states were more frequent (F(95) = 51.14, p = 1.84 × 10^-25^, *BF*_10_ = 2.89 × 10^22^). **B. Computational estimates of perceptual conflict.** Average prediction errors (PE) were elevated during incongruent as compared to congruent perceptual states (F(209) = 158.08, p = 2.29 × 10^-27^, *BF*_10_ = 3.15 × 10^20^). The difference in prediction errors between congruent and incongruent perceptual states was enhanced for higher signal-to-ambiguity ratios (F(209) = 10.41, p = 6.19 × 10^-9^, *BF*_10_ = 2.51 × 10^6^). Overall, prediction errors did not vary across levels of signal-to-ambiguity (F(209) = 0.54, p = 0.75, *BF*_10_ = 0.02). **C. Neural correlates of perceptual conflict.** We found enhanced BOLD responses during incongruent as opposed to congruent perceptual states in right-hemispherical IFC and V5/hMT+, alongside additional clusters in left precentral gyrus, right posterior-medial frontal gyrus (PMF) and right fusiform gyrus (*p_FWE_* < 0.05; displayed for F > 11; see **corresponding Supplementary Table S2**). Importantly, differences in BOLD activity between incongruent and congruent perceptual states were enhanced at higher levels of signal-to-ambiguity in right-hemispheric insula (left panel), inferior frontal gyrus (left and middle panel) and V5/hMT+ (right panel; *pscv* < 0.05 within the main effect of *Congruency;* displayed for F > 4).

Crucially, this pattern was reflected by neural activity in IFC and V5/hMT+: While controlling for variations in BOLD signals associated with reported changes in conscious experience, we observed enhanced BOLD signals during incongruent perceptual states in right-hemispheric IFC and V5/hMT+ (main effect of *Congruency, p_FWE_* < 0.05, Figure 3C; see Supplementary Table 2 for additional activations). As predicted, both right-hemispheric IFC and V5/hMT+ showed larger differences between incongruent and congruent perceptual states at higher signal-to-ambiguity ratios (interaction between *Congruency* and *Signal-to-Ambiguity*, small-volume correction at *psvc* < 0.05 within the main effect of *Congruency*).

This factorial approach to the neural correlates of perceptual conflict was corroborated by model-based fMRI, which provided a complete replication of E1: While controlling for change-related activity, we found that accumulating perceptual prediction errors correlated with neural activity in right-hemispheric IFC and V5/hMT+ (*p_FWE_* < 0.05, Figure 1C and Supplementary Table 1). In comparison to the analysis based on perceptual change events, the dynamic accumulation of perceptual conflict was better at explaining BOLD signals in right-hemispheric IFC (Supplementary Figure S6). Dynamic causal modelling indicated that signals of perceptual conflict were more likely to originate from V5/hMT+ and reached IFC via feedforward effective connectivity (Supplementary Figure S7). Again, the BOLD signal in V5/hMT+ voxels that showed enhanced activity during perception of *leftward* rotation correlated more strongly with perceptual prediction errors when participants experienced *rightward* rotation (*BF*_10_ = 53.36) and vice versa (*BF*_10_ = 8.2; Figure 2B and Supplementary Figure S9B).

Together, E2 confirmed our hypothesis that IFC signals dynamic changes in perceptual conflict that are induced by disambiguating stimulus information. Additional control analyses (Supplementary Materials 9.4) ruled out variations in perceptual uncertainty and temporal imbalances between congruent and incongruent perceptual states as alternative explanations for our fMRI results.

### 5.3 Disruption of neural activity in IFC modulates the dynamics of conscious experience

The independent fMRI experiments E1 and E2 provide converging evidence that IFC detects the conflict inherent in sensory ambiguity. In a third experiment (E3), we asked whether this unconscious detection of perceptual conflict by IFC^29^ is relevant for conscious experience. We reasoned that the signaling of perceptual conflict by IFC might facilitate changes in conscious experience during bistable perception. Consequently, disruption of IFC activity should reduce the frequency of such conflict-driven perceptual changes. To test this hypothesis, we used inhibitory TMS with a theta-burst stimulation protocol^30^ to create *virtual lesions* in IFC.

In E3, we re-invited the participants from E1 for two TMS sessions scheduled on consecutive days. In each session, they first reported changes in conscious experience during two runs of ambiguous structure-from-motion. This was followed by 40 sec of neuronavigated TMS to either IFC or a control location at the cranial vertex (see Methods section 7.3.3 for details). Immediately afterwards, participants again reported their perception during two runs of ambiguous structure-from-motion.

After neural activity in IFC was disrupted by TMS, changes in conscious experience occurred less frequently: For virtual lesions in IFC, we observed prolonged perceptual phase durations (post-pre: 6.86 ± 1.79 sec) relative to the vertex condition (−1.59 ± 2.07 sec, paired t-test: *BF*_10_ = 20.05, Figure 4A). This finding indicates that IFC not only detects gradually accumulating perceptual conflict, but also has a causal role in triggering changes in conscious experience.

**Figure 4.**
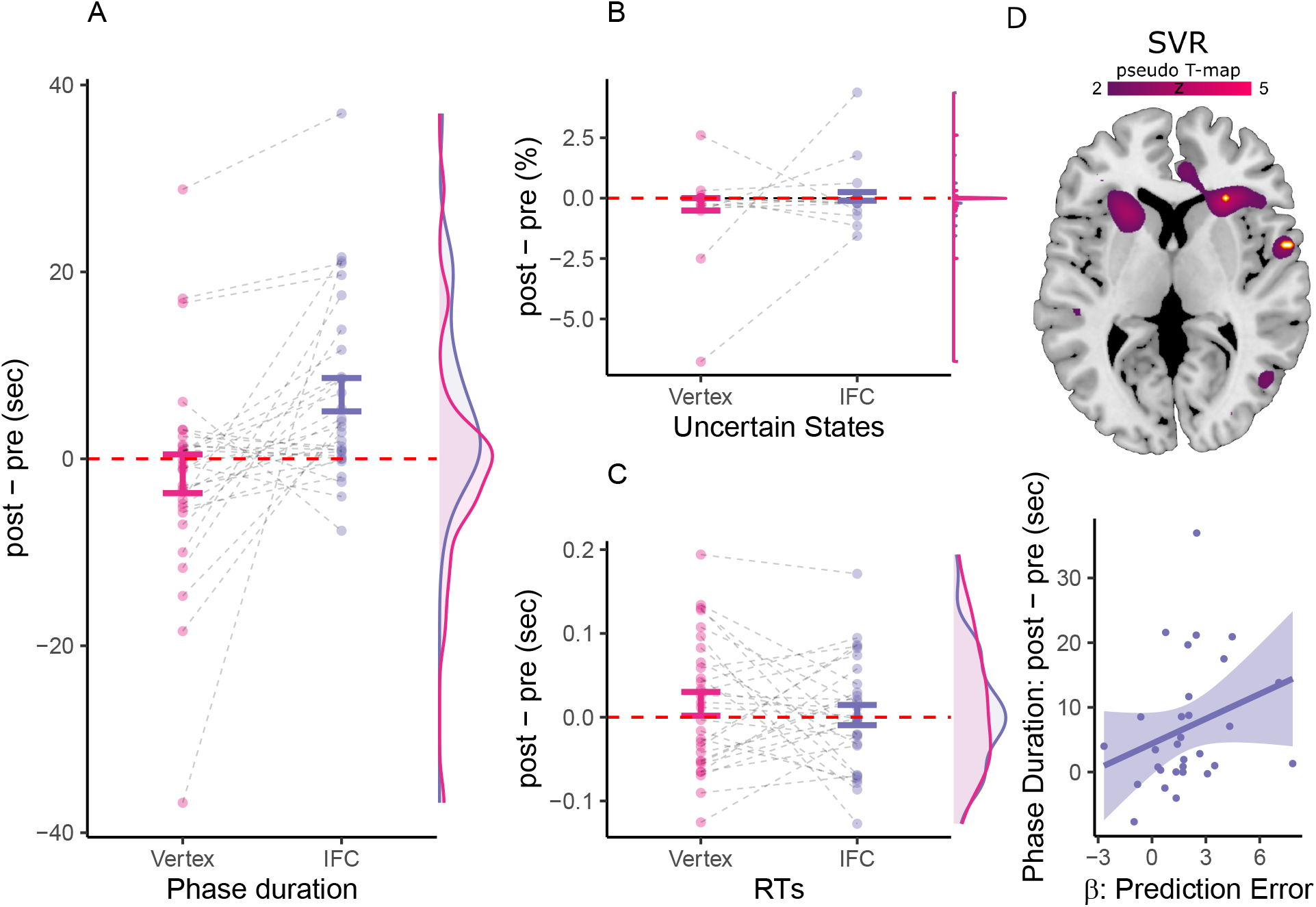
TMS effects on perception. **A. Phase duration.** Virtual IFC-lesions prolonged phase durations relative to the vertex condition (paired t-test: T(29) = −3.44, p = 1.77 × 10^-3^, *BF*_10_ = 20.05) as well as against the baseline recorded prior to IFC stimulation (one-sample t-test: T(29) = 3.85, p = 6.08 × 10^-4^, *BF*_10_ = 51.47). **B. Unclear perceptual states.** TMS to IFC did not alter the frequency of uncertain perceptual states in comparison to the TMS to vertex (T(29) = −1.04, p = 0.31, *BF*_10_ = 0.32). **C. Response times.** Likewise, virtual IFC-lesions did not affect RTs in comparison to control stimulation at vertex (T(29) = 0.77, p = 0.45, *BF*_10_ = 0.26). **D. Representation of perceptual conflict and virtual lesions in IFC.** Whole-brain searchlight decoding (voxels displayed for T > 2, *p_FWE_* < 0.05 highlighted in orange, upper panel) revealed that local fMRI activity patterns in IFC successfully predicted inter-individual differences in the effects of virtual IFC-lesions on conflict-induced changes in conscious experience. Additional clusters were observed in bilateral temporal pole, left posterior-medial frontal gyrus, right superior medial gyrus, right IPL, right V1, right IPL and left middle orbital gyrus. Voxels in hMT+/V5 did not survive FWE-correction across the whole brain. Participants who represented perceptual conflict more strongly in IFC (correlation coefficient *β* of perceptual prediction errors to BOLD signals in individual IFC stimulation sites) showed an enhanced reduction of conflict-induced changes in conscious experience when neural activity in IFC was disrupted by TMS (*ρ* = 0.42, *p* = 0.02; lower panel).

Two additional control analyses addressed alternative accounts of the observed TMS effect on perceptual phase durations. Firstly, previous work has shown that activity in frontal brain regions is elevated at the time of unclear perceptual states during bistability^15^. Here, however, disruption of neural activity in IFC did not alter frequency of unclear perceptual states (post-pre: 0.07 ± 0.18%) in comparison to vertex-stimulation (−0.26 ± 0.26%, *BF*_10_ = 0.32, Figure 4B).

Secondly, when investigating frontal activity as a potential driver of changes in conscious experience during bistable perception, reporting behavior represents a possible confound^7^. “No-report”-paradigms have elegantly addressed this issue, suggesting that a subset of change-related activations in prefrontal cortex may represent the neural correlates of reporting rather than consciousness^16^. Here, we used RTs to ask whether inhibition of activity in IFC impaired the participants’ ability to report on the contents of conscious experience. Changes in RTs did not differ between IFC-(post-pre: 2.55 × 10^-3^ ± 0.01 sec) and vertex-stimulation (0.02 ± 0.01 sec, *BF*_10_ = 0.26, Figure 4C). In Supplementary Materials 9.5-7, we replicate these findings in linear mixed effects modeling (9.5) and rule out exposure effects (9.6) as well as regression toward the mean (9.7) as additional alternative explanations of our results.

In sum, disruption of activity in IFC reduced the frequency of changes in conscious experience during bistable perception. Importantly, we found no evidence for TMS effects on perceptual uncertainty or reporting behavior. These results support the hypothesis that IFC responds to conflicting sensory data by facilitating spontaneous changes in conscious experience, thereby temporarily resolving perceptual conflict^11,14^.

### 5.4 Individual differences in the representation of perceptual conflict predict the effect of virtual IFC-lesions on conscious experience

Finally, we asked whether variability in the neural representation of perceptual conflict could predict inter-individual differences in the effect of virtual IFC-lesions on conscious experience. We used support vector regression to test whether multi-voxel patterns^26^ of conflict-related BOLD activity (E1) contained information on how conscious experience was altered when neural activity in IFC was disrupted (E3). Whole-brain searchlight decoding^31^ revealed that localized multi-voxel BOLD activity in IFC, but not V5/hMT+, predicted the individual effects of virtual IFC-lesions on phase duration (leave-one-out cross-validation with non-parametric permutation testing^32^, *p_FWE_* < 0.05, Figure 4D, upper panel).

In addition, we ensured that neural patterns of conflict representation in IFC selectively predicted the perceptual effects of IFC but not vertex stimulation, and ruled out baseline-differences in phase duration as an alternative explanation of the observed brain-behavior-association (Supplementary Materials 9.8). Univariate analyses confirmed that virtual IFC-lesions reduced the frequency of changes in conscious experience to a greater extent in participants who represented perceptual prediction errors more reliably at IFC stimulation sites (Figure 4D, lower panel).

At the level of IFC, inter-individual differences in detecting conflicting sensory information were thus directly linked to how strongly prefrontal brain activity impacted on conscious experience, closing the loop between feedforward and feedback processing of perceptual conflicts.

## 6 Discussion

In this work, we found compelling evidence for a active role of IFC in conscious experience: Two independent fMRI experiments demonstrated that IFC signals the conflict that emerges between conscious experience and the underlying sensory data. Crucially, TMS-induced virtual lesions revealed that IFC facilitates changes in conscious experience that occur in response to accumulating perceptual conflict.

### 6.1 IFC detects and resolves perceptual conflict during bistable perception

At first glance, our results may seem at odds with the well-established dynamic system account of bistable perception^33^. This view proposes that, in the context of conflicting stimulus information, changes in conscious experience result from local mechanisms such as inhibition, adaption or noise^8^. Along these lines, neural activity occurring within sensory corteces may be sufficient to distill unambiguous conscious experiences from conflicting sensory data.

Indeed, our data verify that the contents of conscious experience can be decoded from BOLD-activity at the level of V5/hMT+ (Supplementary Figure S8). Concurrently, we found that V5/hMT+ generates signals of accumulating perceptual conflict that originate from voxels coding for the currently suppressed stimulus interpretation (Figure 1 and 2). In the suppressed voxels, BOLD signals progressively increase prior to changes in conscious experience. In mechanistic terms, these escalating signals of perceptual conflict may be generated by neural populations that gradually escape from inhibition, as adaption reduces the activity in competing neural populations that represent the currently dominant stimulus interpretation. Our results therefore do not contradict the dynamic system account of bistable perception, but suggest that the *implementational* concept of local adaption-and-inhibition^33^ and the *algorithmic* hypothesis of dynamic conflict-accumulation^11,14^ are, in fact, complementary^8^.

Importantly, however, our results clearly indicate that the processing of perceptual conflicts does not end at the level of sensory brain regions, but reaches prefrontal cortex through feedforward processing from V5/hMT+ to IFC (Figure 1). Crucially, we found that disrupting neural activity in IFC reduces the impact of perceptual conflict on conscious experience (Figure 4A and D). This indicates that IFC activity is not just a downstream consequence of perceptual events that are realized within hMT+/V5, but actively contributes to the resolution of sensory ambiguity via feedback processes. Together, our findings thus reconcile the feedforward- and feedback-accounts of bistable perception^8^, suggesting a hybrid computational function of IFC in conscious experience: the detection and resolution of perceptual conflict.

Such a hybrid model^12^ not only aligns with previous work suggesting a causal influence of prefrontal feedback on bistable perception^8^, but also provides a plausible explanation for the absence of prefrontal activity when perceptual events remain invisible^17,18^: Possibly, the capacity of IFC to detect perceptual conflict through feedforward processing may be limited to situations in which the competing states are perceptually distinguishable. When they are not, IFC may fail to read out conflicting stimulus representations^23^, leaving the resolution of perceptual conflict to sensory brain regions^34^. By analogy, our results account for the increase in neural activity observed during unclear or *mixed* conscious experience^15^, since such perceptual states represent instances of enhanced perceptual conflict and are typically linked to perceptual changes. Future experiments could test this prediction by correlating conflict-related BOLD-activity in V5/hMT+ and IFC with a parametric modulation of the salience evoked by perceptual events during bistable perception.

### 6.2 Attention, response behavior and cognitive control as alternative accounts for IFC’s role in bistable perception

IFC has been implicated in various domains of cognition, including attention^35,36^, response behavior^17^ and cognitive control^37^. IFC may therefore exert its influence on conscious experience through one of these alternative cognitive functions, rather than participating directly in the resolution of perceptual conflicts.

First, neural activity prefrontal cortex is known to support sustained attention^35^. During bistable perception, changes in conscious experience occur less frequently when attention is withdrawn^38^. One may therefore argue that virtual IFC-lesions may have impaired the participants’ ability to attend to the experimental task and, consequently, reduced the frequency of perceptual changes. Yet, two observations argue against this proposition: Firstly, we did not observe any effect of virtual IFC-lesions on response times (Figure 4C), which closely link to levels of on-task attention^39^. Secondly, support vector regression revealed that the prefrontal impact on conscious experience is specifically predicted by how strongly IFC activity tracked the accumulation of perceptual conflict (Figure 4D). Sustained attention, in turn, is unlikely to increase systematically over the course of each perceptual phase. It is therefore improbable that the prolongation of perceptual phase durations following virtual IFC-lesions can be explained solely on the ground of a global reduction in sustained attention. To directly test this caveat, future work could combine virtual IFC-lesions with a parametric modulation of on-task attention during bistable perception.

Second, it has repeatedly been proposed that prefrontal cortex supports only the down-stream report of changes of conscious experience that are realized at earlier processing stages^16,17^. Yet, a selective impairment of motor behavior seems unable to explain why conflict-induced change in conscious experience are less likely to occurr after virtual IFC-lesions (Figure 4A), which left response times unaltered. In addition, our fMRI analyses reveal consistent correlations between IFC activity and accumulating perceptual conflicts while explicitly controlling for the neural correlates of actively reported changes in conscious experience (Figure 1). Based on these findings, we conclude that the often-reported finding of change-related IFC activity is in fact likely to reflect the peak of accumulating perceptual conflict instead of the reported event per se (Supplementary Figure S4–6). This aligns with recent results showing that change-related prefrontal activity seems to persist in no-report paradigms of bistable perception^40^. To further substantiate the view that IFC activity is not primarily linked to motor behavior, future experiments should test whether the prefrontal representation of perceptual conflict and its causal effect on conscious experience are modulated by active report.

Third, the gradual accumulation of IFC activity toward a toward changes in conscious experience during bistable perception may alternatively be explained by processes related to cognitive control^37^ and the anticipation of future events^41^: As the perceptual phase grows longer, participants may become increasingly prone to expect a change in perception. Conversely, they may be more relaxed once an event has occurred. Since average phase durations are quite consistent within individuals (Supplementary Figure S2), participants may be capable of predicting the approximate timing of changes in conscious experience during bistable perception. Thus, phasic changes in the anticipation of upcoming events may indeed be compatible with the dynamic changes of BOLD observed in IFC.

It may be speculated that, when anticipating a perceptual event, participants could try to voluntarily increase the likelihood of a change in conscious experience^42^. Virtual lesions in dorsolateral prefrontal cortex (DLPFC) have been shown to impair the capacity to exert voluntary control over ambiguous structure-from-motion stimuli^43^. Under this assumption, the observed effect of virtual IFC-lesions on conscious experience could be mediated via an impairment of cognitive control, rather than via a mechanism that resolves perceptual conflicts.

In our study, however, participants were naive to the ambiguity in the visual display. Moreo-ever, they were explicitly instructed to passively view the display and report their conscious experience of the stimulus. In contrast to de Graaf et al., who found an effect of prefrontal TMS only on the voluntary control of bistable perception, we observed clear evidence for a prolongation of phase duration during passive viewing (Figure 4). Next to differences in sample size (N = 30 vs. N = 10^43^), this discrepancy may also be due to the target region: While we stimulated IFC and defined stimulation sites based on the neural correlates of perceptual conflict in each participant individually, de Graaf et al. stimulated DLPFC using standard 10/20 electroencephalography coordinates (F4). Yet, to fully resolve the question whether anticipation induces prefrontal mechanisms of cognitive control that represent an additional driving factor for spontaneous perceptual changes, future work should use disambiguated stimuli to induce specific temporal expectations and test their effect on conscious experience during bistable perception.

### 6.3 IFC regulates the access of conflicting information into conscious experience

With respect to the role of prefrontal cortex in consciousness, our results speak against the notion that IFC activates merely as a consequence of perceptual events that are generated within sensory cortices^15–17^. As a significant extension, our work associates IFC with a specific computational function for conscious experience: In iterative feedback- and feedforward-interactions with sensory brain regions, IFC may determine how swiftly conscious experience is updated in response to perceptual conflict^11,12,14^. Intriguingly, this finding aligns with recent neural recordings in monkeys suggesting that prefrontal state fluctuations precede changes in perception during no-report binocular rivalry^40^.

By controlling the entry of conflicting information into consciousness, IFC may ensure that perception is altered when discrepancies between conscious experience and sensory data have accumulated over time, but remains stable when perceptual conflicts are transient. In mechanistic terms, feedback from IFC to sensory cortex could support this function by decreasing the mutual inhibition between competing neural populations^44^, by increasing the rate of adaption^45^ or by upregulating the level of noise in perceptual processing^46^. In these non-exclusive scenarios, feedforward-feedback-loops between sensory and prefrontal cortex could benefit perception by facilitating changes in the content of conscious experience only in situations of escalating perceptual conflict.

Beyond the context of regulating the access of conflicting sensory information into conscious experience, IFC may play a similar adaptive role in orienting toward relevant stimuli^35^, in detecting change^47^, or in allocating object-based attention^36^. Altered states of consciousness such as hallucinations^48^ could therefore relate directly to an impaired processing of sensory information in IFC. Indeed, previous research has associated sensitivity to perceptual conflict with the severity of hallucinations^28^. Correspondingly, functional imaging has repeatedly linked hallucinations to neural activity in IFC^49,50^. Non-invasive brain stimulation of IFC may thus represent a promising new approach in the search for the therpeutic modulation of altered states of consciousness.

While our results demonstrate that IFC is relevant for transforming ambiguous sensory information into unambiguous conscious experiences, they do not provide clear evidence for a representation of perceptual content in prefrontal cortex as a whole (Supplementary Figure S8A). Moreover, the observed trend-wise decoding of specific conscious experiences in IFC (Supplementary Figure S8B) has to be interpreted with caution, since our approach did not dissociate perceptual contents from behavioral reports. Yet, previous research has repeatedly shown that prefrontal cortex may indeed represent the contents of conscious experience^51,52^. Intersecting computational models of dynamic conflict-accumulation^14^ with no-report paradigms of bistable perception will enable future research to test whether the contents of conscious experience are represented^53^ or multiplexed^54^ within the neural correlates of perceptual conflict, creating exciting new opportunities to better understand the role of prefrontal cortex in consciousness.

## 7 Methods and Materials

### 7.1 Participants

Experiment E1 consisted of a behavioral pre-test (Runs 1 and 2) and an fMRI-experiment (Runs 3 - 5). We recruited a total of 35 participants. Based on the behavioral pre-test, we excluded two participants who performed at chance-level when discriminating the direction of rotation of a fully disambiguated structure-from-motion stimulus. Thus, 33 participants took part in the fMRI-experiment (21 female, mean age: 27.3 ± 1.42 years). All participants agreed to be contacted later for a follow-up experiment using TMS (E3, see below).

Experiment E2 consisted of a behavioral pre-test and a fMRI-experiment. We recruited a total of 23 participants. We excluded three participants who performed at chance-level when discriminating the direction of rotation of a fully disambiguated structure-from-motion stimulus. The final sample thus consisted in 20 participants (11 female, mean age: 27.7 ± 0.98 years).

For experiment E3, we re-invited the participants from E1 to two TMS session scheduled on consecutive days. From this group, one participant could not be re-contacted at the time of the TMS-experiment. Two further participants did not tolerate the TMS-procedure. The final TMS-sample thus consisted of 30 participants (19 female, mean age: 27.33 ± 1.56 years).

All participants were right-handed, showed corrected-to-normal vision, had no prior neurological or psychiatric medical history and gave written informed consent prior to taking part in the study. All procedures were approved by the ethics committee at Charité Berlin.

### 7.2 Stimuli

Stimuli were presented using Psychtoolbox 3^55^ and Matlab R2007b (behavioral pre-test: CRT-Monitor at 60 Hz, 1042 x 768 pixels, 60 cm viewing distance and 30.28 pixels per degree visual angle; fMRI: LCD-Monitor at 60 Hz, 1280 x 1024 pixels, 160 cm viewing distance and 90.96 pixel per degree visual angle; TMS: LCD-Monitor at 60 Hz, 1280 x 1024 pixels, 60 cm viewing distance and 37.82 pixels per degree visual angle).

#### 7.2.1 Random dot kinematograms

Throughout E1, E2 am E3, participants indicated their perception of a discontinuous random-dot kinematogram (RDK, see Supplementary Video 1). In this stimulus, random dots distributed on two intersecting rings induce the perception of a spherical object rotating either left- or rightward around a vertical axis^19^ (diameter: 15.86°, rotational speed: 12 sec per rotation, rotations per block: 10, individual dot size: 0.12°). Each run consisted of six blocks of visual stimulation (120 sec), separated by fixation intervals (behavioral pre-tests: 5 sec; fMRI- and TMS-experiments: 10 sec).

Depending on the experimental condition (see Supplementary Figure S1), the RDK could appear in three configurations: Complete ambiguity, levels of graded ambiguity and complete disambiguation. Complete ambiguity was achieved by presenting identical stimuli to the two eyes. This induced periodic changes in conscious experience (also dubbed *endogenous transitions*) between left- and rightward rotation (i.e., bistable perception).

For complete disambiguation, we used red-blue filter glasses (left eye: red channel, right eye: blue channel) to attach a stereo-disparity signal (1.8° visual angle) to all dots on the stimulus surface. By inverting the direction of rotation, we created stimulus-driven or *exogenous* changes in conscious experience.

During graded ambiguity^28^, we varied the proportion of disambiguated stimulus dots between 15%, 30%, 45%, 60%, 75% and 100% (conditions D1 to D6). This variation in the signal-to-ambiguity ratio parametrically modulated the perceptual conflict between conscious experience and visual stimulation: We predicted that perceptual conflict (and associated neural activity) should be enhanced during incongruent as compared to congruent perceptual states. Furthermore, this enhancement should increase at higher signal-to-ambiguity ratios. During runs with graded ambiguity, conditions D1 to D6 appeared in random order. Within each block, we introduced random changes in the direction of disambiguation (i.e., whether the parametric 3D cues enforced rightward or leftward rotation). The individual frequency of exogenous stimulus changes during graded ambiguity was determined based on the frequency of conflict-induced changes in conscious experience during full ambiguity.

Importantly, participants were naive to the potential ambiguity in the visual display and explicitly instructed to passively experience the stimulus, reporting their perception via button-presses (right index-finger: rotation of the front-surface to the left; right ring-finger: rotation to the right; right middle-finger: unclear direction of rotation) on a USB keyboard or a MRI-compatible button-box, respectively.

We based our behavioral analyses on perceptual events as reported by the participants. Since the RDK is not depth-symmetric over all rotational angles^19,56^, changes in conscious experience occurred only at overlapping configuration of the stimulus (see Supplementary Figure S2A-B). We thus corrected the timing of perceptual events to the last overlapping configuration of the stimulus preceding the button-press, representing the perceptual time-course as a discrete sequence of perceptual states (rotation of the front-surface to the right/left and unclear direction of rotation).

To describe the temporal dynamics of bistable perception, we computed *average phase durations* (the time spent between two changes in conscious experience, i.e., multiples of the 1.5 sec inter-overlap-interval). The content of conscious experience was reflected by the dependent variables *directed bias* (the percentage of rightward perceptual states relative to the total number of perceptual states), *absolute bias* (the absolute difference between the absolute bias and chance level at 50%) and the percentage of *uncertain states*. To characterize processes involved in the behavioral report of perception, we computed *response times* by subtracting the timing of the last preceding overlapping configuration from the timing of the behavioral responses indicating a perceptual event. The impact of sensory data on perception was depicted by the dependent variable *perceptual congruency* (percentage of perceptual states congruent with the disambiguating 3D signal).

#### 7.2.2 Heterochromatic flicker photometry

When using filter glasses (Experiment E2), the perceived direction rotation of RDKs can be biased by differences in the subjective luminance between red and blue (Pulfrich effect^57^). To estimate subjective equiluminance, we presented red and blue circles (diameter: 6.45°) alternating at a frequency of 15 Hz. Differences in subjective luminance of red and blue stimuli led to the experience of flicker. Participants reduced the flicker by adjusting the luminance of the red stimulus initially presented at a random luminance between 0% and 255% relative to the blue stimulus presented at a fixed luminance. Average equiluminance estimated across 10 such trials determined the monitor- and participant-specific luminance of the red- and blue-channels (average blue-channel luminance relative to red-channel: 2.02 ± 0.09).

#### 7.2.3 2D control stimuli

At higher levels of signal-to-ambiguity, perceptual states were less likely to be incongruent with the disambiguating stimulus information. Hence, increments in the signal-to-ambiguity ratio increased the temporal imbalance between congruent and incongruent perceptual states. To test for potential confounds introduced by temporal imbalance, we constructed a 2D control version (E2, Run 8) of the bistable RDK (identical stimulus diameter, number, size and speed of dots). Participants reported the direction of planar, horizontal 2D motion. For each participant, changes in the direction of planar dot motion were determined by both the temporal imbalance between congruent and incongruent perceptual phases (separately for conditions D1-D6) as well as the average frequency of changes in conscious experience observed in the main experiment (E2, Runs 5-7). We randomized the motion direction associated with reduced presentation-time.

### 7.3 FMRI

#### 7.3.1 Acquisition and preprocessing

For E1 and E2, we recorded a T1-weighted MPRAGE sequence (voxel size 1 x 1 x 1 mm) for anatomical images and used T2-weighted gradient-echo planar imaging (TR 2000 ms, TE 25 ms, voxel-size 3 x 3 x 3 mm) to obtain a total of 400 BOLD images per run on a Siemens Prisma 3-Tesla-MRI-system (64-channel coil). Our pre-processing routine was carried out within SPM12 and consisted in slice time correction with reference to the middle slice, standard realignment, coregistration and normalization to MNI stereotactic space using unified segmentation. For standard analyses and support vector regression^32^, we applied spatial smoothing with 8 mm full-width at a half-maximum isotropic Gaussian kernel. For the analysis of voxel biases and support vector classification, we used unsmoothed data.

#### 7.3.2 General linear models

To test for the neural correlates of perceptual conflict during sensory ambiguity in E1 and E2, we extracted dynamic perceptual prediction errors from the predictive-coding (PC) model of bistable perception^14^, which was inverted based on behavioral data. This *model-based* fMRI approach (**GLM-PC**) defined visual stimulation by stick-regressors aligned to the overlapping configurations of the structure-from-motion stimulus (*overlaps*). Relative to the *overlaps*, we defined two parametric regressors ordered as follows: (1) perceptual *changes* (binary; 0: no change, 1: change) and (2) absolute *prediction errors* (continuous, ranging from 0 to 1). For the analysis of direction-specific effects in voxel biases within V5/hMT+, the overlaps and the associated parametric modulators were modeled separately according to the current perceptual state (left-vs-rightward rotation). In addition to standard GLMs, we performed Bayesian second-level statistics^22^ to compare the explanatory power between change-related models and prediction-error related models with regard to BOLD activity in IFC. To this end, we deleted one of the two parametric modulators in GLM-PC, creating Log-Evidence-Maps for the two degraded models (“PE only” vs. “Change only”; z-scored parametric modulators).

In E2, we used an additional GLM (**GLM-Congruency**) to analyze BOLD activity during graded ambiguity independently of the assumptions inherent in the predictive-coding model of bistable perception. Next to perceptual changes (*T*, stick-function), this GLM represented perceptual states by box-car regressors defined according to two factors: Firstly, perceptual states could be congruent (*C1*) or incongruent (*C2*) to conscious experience. Secondly, visual stimulation varied across six levels of signal-to-ambiguity (*D1-D6*). The GLM’s design matrix contained all combinations between the two factors ([C1D1 C1D2 (…) C1D6 C2D1 C2D2 (…) C2D6 T]). By analogy, we tested for a potential effect of temporal imbalances between congruent and incongruent perceptual phases in the fMRI control-experiment (run E2(8)). **GLM-Control** defined prolonged (*A1*) and shortened (*A2*) perceptual phases separately for all levels of temporal imbalance (Levels *I1* to *I6*; design-matrix = [A1I1 A1I2 (…) A1I6 A2I1 A2I2 (…) A2I6 T]).

In E3, we identified individual IFC stimulation sites based on the fMRI data acquired in E1. To delineate IFC independently of the assumptions inherent in the predictive-coding model of bistable perception (see GLM-PC), we adopted the conventional change-related fMRI approach to IFC^15,17,56^, representing endogenous perceptual changes as stick-functions and visual stimulation as a box-car regressor (**GLM-Changes**).

In all GLMs, we convolved the outlined regressors with the canonical hemodynamic response function (SPM12), added six rigid-body realignment parameters as nuisance covariates, applied high-pass filtering at 1/128 Hz and computed first-level one-sample t-tests against baseline. On the second-level, the resulting images were submitted to second-level one-sample t-tests (GLM-PC) or full factorial models (GLM-Congruency and GLM-Control).

Second-level results were thresholded at p < 0.05 (FWE-corrected across the whole brain; SVC within orthogonal activation maps for GLM-Congruency). For Bayesian second-level statistics^22^, we display second-level results at an exceedance probability of 95% for “PE only”.

#### 7.3.3 Stimulation sites at IFC and vertex

In E3, we defined individual IFC coordinates for neuronavigated TMS based on BOLD activity associated with perceptual changes (data acquired during E1). Using GLM-Changes, we identified the peak voxel for “Changes vs. baseline” (first-level contrast at p < 0.005, uncorrected) within a literature-based IFC search-sphere (radius = 5 mm; centre = [57 17 10]). This location was motivated by the neural correlates of conflict-driven as opposed to stimulus-driven changes in conscious experience in a closely related structure-from-motion stimulus^14^. Across participants, average stimulation sites were located at MNI = [55.6 ± 0.4 15.5 ± 0.39 10 ± 0.49].

By informing the TMS-intervention based on the conventional change-related approach to IFC^15,17^, we delineated the IFC stimulation site independently of our computational model of bistable perception^14^. As shown above, change-related activity coincided with the neural correlates of perceptual prediction errors (Figure 1C, Supplementary Figure S5B), which had more explanatory power with regard to BOLD signals in IFC^14^ (Supplementary Figure S6). As expected, activity in the IFC stimulation site was thus highly correlated to perceptual prediction errors (average regression coefficients in spherical ROIs of 10 mm radius around individual coordinates: *β* = 1.79 ± 0.38; T(32) = 4.75, p = 4.15 × 10^-5^, *BF*_10_ = 565.3).

The control stimulation site at vertex was determined by anatomical (T1) scans (MNI = [0 −25 85]). Given the spatial resolution of neuronavigated TMS^58^, vertex-TMS was extremely unlikely to exert local effects on any additional neural correlates of bistable perception (Supplementary Table 1).

#### 7.3.4 Regions-of-interest (ROI)

All ROIs were defined independently of the computational model of bistable perception^14^ outlined in Supplementary Materials 9.2. With respect to IFC, we defined spherical ROIs (radius: 10 mm) around the individual IFC-TMS coordinates (see above). To delineate V5/hMT+, we constructed a search sphere (radius: 5 mm) around the peak-voxel for the second-level contrast “Visual Stimulation vs. baseline” (GLM-Changes, *p_FWE_* < 0.05) within an anatomical mask for V5/hMT+^59^. Based on this search sphere, we constructed individual V5/hMT+-ROIs (radius: 10 mm) centered around the individual peak coordinates of the corresponding first-level contrast (p < 0.005, uncorrected). Within these ROIs, we defined voxels with biases for righward- and leftward perceptual states (L- and R-population) by thresholding the contrasts for “left vs. rightward perceptual states” and vice versa at a T-value of 1 (GLM-PC). Functional ROI-based analyses were carried out in Marsbar (http://marsbar.sourceforge.net/).

#### 7.3.5 Anatomic Labeling

All anatomic labels were obtained from the Anatomy Toolbox^59^. The IFC was defined by the combination of anterior insula and inferior frontal gyrus (past triangularis and pars opercularis).

### 7.4 TMS

In E3, we used TMS with a theta-burst protocol to induce virtual lesions in the two stimulation sites (i.e., the target-region in right-hemispherical IFC and the control-region at the cranial vertex, see above). TMS was delivered in two separate TMS-sessions on two consecutive days. We counterbalanced the order of IFC-vs vertex-TMS across participants. Participants performed two runs of the experiment prior to TMS and two runs immediately after TMS.

We delivered TMS with a focal, figure-of-eight-shaped coil equipped with active cooling. Pulses where generated using a standard MagPro R30 stimulator (MagVenture Ltd, Farum, Denmark). Stimulation was guided by online neuro-navigation based on individual target regions projected onto the participants’ T1 scans using the Localite TMS Navigator (Localite GmbH, Bonn, Germany) with an optical tracking camera PolarisVicra (Northern Digital Inc., Ontario, Canada).

The coil was positioned tangentially to the subjects’ head and adjusted such that the electric current in the center of the coil would run perpendicular to the course of the inferior frontal sulcus. Prior to each session, we identified individual resting motor thresholds (rMTs) for the right first dorsal musculus interosseous (FDI) by stimulation of left-hemispherical motor cortex^60^. The coil was held tangentially to the subject’s skull at a 45° angle to the parasagittal line (4 cm lateral and 1 cm anterior to the vertex). The search for the hot-spot was additionally guided through the optical tracking system in order to locate the handknob. In order to find the rMT hot-spot, we started with 55% Maximum Simulator Output (MSO) and increased the intensity in 5% steps. If a motor evoked potential (MEPs) was elicited, adjustments were made in 1% steps. Pulses for MT search were delivered with a minimum of 5 sec delay in order to avoid any change in excitability due to repeated stimulation. MEPs were recorded from the right FDI using self-adhesive gel electrodes in a standard belly-tendon fashion. RMTs were defined as the percentage of maximum stimulator output needed to evoke 50μV MEPs peak-to-peak in five out of ten consecutive trials (average rMT in vertex sessions: 41.67 ± 1.12% MSO; IFC sessions: 40.9 ± 1.15% MSO; paired t-test: T(29) = 0.89, p = 0.38, *BF*_10_ = 0.28).

The theta-burst TMS-protocol consisted in a total 600 pulses applied within 40 sec (50-Hz bursts with three pulses applied in intervals of 200 ms) at an intensity of 80% rMT. Stimulation parameters were in line with published safety guidelines and were chosen to produce a decrease in cortical excitability^30,61–63^ throughout the 25 min test-phase following TMS.

### 7.5 Brain-behavior associations

To relate inter-individual differences in the representation of perceptual conflict to the effects of virtual IFC-lesions on conscious experience, we assessed brain-behavior associations in both a *univariate* and a *multivariate* approach. In the *univariate* approach, we conducted a standard ROI-based analysis, extracting individual regression coefficients *β* of perceptual prediction errors to BOLD signals from individual IFC stimulation sites. We then used Spearman correlation to test whether individual *β* estimates predicted the behavioral effects of virtual lesions.

In *multivariate* pattern analysis, we predicted the effects of virtual IFC-lesions based on localized pattern of BOLD-activity measured across the whole brain. Using searchlight decoding^31^, we extracted multidimensional pattern vectors from spherical clusters (8 mm radius) centered around each voxel within the individual participants’ T-maps for *Perceptual prediction error vs. baseline* (GLM-PC). These multidimensional vectors thus reflected how locally distributed patterns of fMRI activity represented perceptual prediction errors.

Based on these multidimensional vectors, we trained a support vector regression machine (SVR; linear kernel, constant regularization parameter of 1; implemented in LIBSVM, http://www.csie.ntu.edu.tw/~cjlin/libsvm) to predict the individual participants’ post-pre difference in phase duration associated with virtual lesions in IFC. At each voxel, we performed 30 iterations of leave-one-out cross-validation, using the labeled data for 29 out of the 30 participants as the training-set and the remaining participant’s data for testing. Prior to training, we normalized both the continuous labels and the multidimensional pattern vectors (i.e., *x_norm_* = (*x* – *min*(*x*)) / (*max*(*x*) – *min*(*x*)), with normalization parameters derived from the training set alone^32^.

In the test-set, we assessed predictive performance by calculating Pearson’s correlation coefficients between the actual and the predicted difference in phase duration associated with virtual lesions in IFC. P-values were computed at each voxel using nonparametric permutation testing. To create a null-distribution of correlation coefficients at each searchlight voxel, we repeatedly trained and tested the SVR with randomly permuted labels.

We considered prediction accuracy to be significant if permutation testing indicated that the probability of the true correlation occurred at *p_FWE_* < 0.05. Prediction accuracy was thus assessed with Bonferroni-correction for multiple comparisons across all voxels in the whole volume of the brain. Therefore, the boundary p-value surving FWE-correction was defined by p < 0.05/n, with n = 48 833 voxels inside the whole-brain volume. Permutation testing thus required up to 1/(0.05/n) = ~9770000 iterations at each voxel. We reduced the computational load by aborting permutation testing for a voxel where three values of the test statistic exceeded the true correlation coefficient^32^.

For visualization (Figure 4D), we computed *pseudo T-values* by drawing T-values corresponding to the nonparametric p-values from an inverted student’s T-distribution. We smoothed the resulting T-map with an 8 mm Gaussian kernel.

## 9 Supplementary Materials

### 9.1 Stimulus characteristics

In this work, we elicited bistable perception by presenting a discontinuous structure-from-motion stimulus (Supplementary Video S1). Experiments E1 and E3 used fully ambiguous versions of the stimulus and were conducted in the same sample of participants. In E2, the stimulus was either fully ambiguous or parametrically disambiguated (*graded ambiguity*). In each study sample, we also collected data for one run of fully disambiguated structure-from-motion stimuli alternating at a probability of 15% per overlapping configuration. These control runs allowed us to assess (i) whether participants followed the experimental instructions during ambiguity and (ii) whether there was a difference in the frequency of unclear perceptual states between fully ambiguous and fully disambiguated stimuli. Here, we assess the characteristics of the structure-from-motion stimulus across experiments E1, E2 and E3 (see Main Text for data on *phase duration* and *RTs*).

Individual participants showed small but significant *biases* toward one of the two perceptual alternatives: The more prevalent alternative accounted for 64.87 ± 1.07% of perceptual states (*absolute bias*, T(54) = 13.88, p = 1.9 × 10^-19^, *BF*_10_ = 2.66 × 10^16^; one-sample t-test). Across participants, however, none of the two perceptual alternatives was preferred over the other (*directed bias*): 48.47 ± 1.01% of perceptual states were assigned to rightward rotation of the sphere (T(54) = −1.5, p = 0.14, *BF*_10_ = 0.42).

*Unclear perceptual states* were rare (0.6 ± 0.18%) and more frequent in comparison to a disambiguated control stimulus (0.04 ± 0.03%; T(54) = 3.13, p = 2.81 × 10^-3^, *BF*_10_ = 11; paired t-test). Accuracy of perceptual reports for this disambiguated stimulus was high (96.75 ± 1.42%), indicating that participants followed the experimental instructions.

### 9.2 The predictive-coding model of bistable perception

In this work, we investigated how neural activity in IFC related to the perceptual conflict inherent in ambiguous sensory information. Next to a *standard* assessment of perceptual conflict (see GLM-Congruency, E2), we applied an established *computational model* of bistable perception^14,28,64^. By inverting this model, we estimated perceptual prediction errors as a quantitative representation of perceptual conflict.

In this supplement, we provide a mathematical description of the computational model of bistable perception. In addition, we describe how the model was inverted based on behavioral data. In simulation analyses, we illustrate the relation between model parameters (*π_IPS_*: the initial belief in the stability of the visual environment; *π_ERROR_*: the impact of perceptual conflict on the belief in the stability of the visual environment; *π_DIS_*: the participants’ sensitivity to disambiguating stimulus information) and the temporal characteristics of conscious experience *y*. With this, we derive quantitative predictions for the behavioral and imaging analyses outlined in the Results section.

#### 9.2.1 Model description

Throughout the experiments E1 to E3, we presented a rotating discontinuous structure-from-motion stimulus. Participants reported whether they perceived the front surface of the object as rotating to the left or right. During full ambiguity, the direction of rotation spontaneously changed at a specific frequency (phase duration) in each participant. During graded ambiguity^28^, we experimentally manipulated the stimulus by introducing *disambiguating stimulus information* in form of 3D cues. Depending on the signal-to-ambiguity ratio, this disambiguating stimulus information biased conscious experience toward stimulus-congruent perceptual states.

Here, we explain how sensory data and implicit beliefs about the stability of the sensory environment give rise to perceptual states *y* during full and graded ambiguity. We adopt a Bayesian approach assuming that perceptual states are determined by *posterior probability distributions*. Posterior probability distributions result from the combination of currently available sensory data (the *likelihood distribution*) with information acquired from previous visual experience (the *prior distribution*).

During full ambiguity, our model assumes a bi-modal likelihood distribution representing balanced evidence for both perceptual interpretations. Graded ambiguity shifts the balance of the likelihood in the direction of one perceptual interpretation at the expense of the other. In this context, sensory information is described by the direction of disambiguation (*μ_DIS_*) and the amount of disambiguation (i.e., the *signal-to-ambiguity* ratio; defined for the condition D1-D6 of experiment E2). As a free parameter, *π_DIS_* reflects the individual impact of disambiguating stimulus information on conscious experience. This is equivalent to the participants’ sensitivity to disambiguating stimulus information during graded ambiguity.

The prior, in turn, is modeled as a uni-modal distribution centered on the previously dominant perceptual interpretation. It acts as an implicit belief in the stability of the environment. The prior is defined by the current perceptual state (*μ_stability_*) and its impact on future conscious experience (*π_stability_*). Two free parameters define the the temporal evolution of *π_stability_*: *π_IPS_* reflects the maximum value of *π_stability_*, which we allocate to the beginning of a perceptual phase. In addition, we assume that *π_stability_* decays linearly during a perceptual phase. This linear decay (with a lower bound at 0) occurs relative to the impact (or *precision*) of perceptual prediction errors (*π_ERROR_*, see below).

The model combines bimodal likelihood and unimodal stability prior. This computes the available evidence for both interpretations of the sensory data. Crucially, once a percept is established, the residual evidence for the suppressed perceptual state constitutes a perceptual prediction error. Relative to the precision of the prediction error (*π_ERROR_*), this quantitative representation of perceptual conflict leads to a linear reduction in the precision of the stability prior. Over time, this results in escalating prediction errors and a dynamic shift of the posterior distribution toward the currently suppressed perceptual interpretation. This entails an increasing probability of a change in conscious experience. Once the change has occurred, the stability prior shifts to the now-dominant stimulus interpretation and its precision is re-set to an initial value (*π_IPS_*). As predicted by predictive-coding theories of perceptual inference^11,65^, prediction errors are thus minimized after the observer adopts a new perceptual interpretation.

In addition, our model assumes a modulation of prediction error accumulation by disambiguating stimulus information: When the current perceptual state is congruent with the disambiguating sensory evidence, our model predicts that prediction errors are reduced relative to full sensory ambiguity. Conversely, when perception is incongruent with the disambiguating sensory evidence, our model assumes enhanced perceptual prediction errors. Importantly, the strength of this enhancement/reduction in prediction errors scales with the amount of sensory evidence during graded ambiguity (i.e., the *signal-to-ambiguity* ratio) and the participants’ sensitivity to disambiguating stimulus evidence (*π_DIS_*).

Hence, three free parameters control the perceptual dynamics produced by our model: The intial precision of the stability prior *π_IPS_*, the precision of perceptual prediction errors *π_ERROR_*, which governs the rate of linear decay in the precision of the stability prior over time, and, in the case of graded ambiguity, the participants’ sensitivity to disambiguating stimulus evidence *π_DIS_*. We infer these parameters by inverting our model based on the sequence of percepts *y* indicated by the participants and, in the case of graded ambiguity, the available sensory information (*μ_DIS_*: direction of disambiguation; *SAR*: signal-to-ambiguity ratio)

Since changes in conscious experience for non-depth-symmetrical structure-from-motion stimuli occur exclusively at overlapping stimulus configurations^19,56^, we represent percepts and all further model quantities in discrete time points *t* defined by stimulus overlaps. For computational expediency, our model assumes Gaussian probability distributions defined by mean and precision (inverse of variance).

At each timepoint *t*, we compute the probability of the two percepts based on the posterior distribution *P*(*θ*):

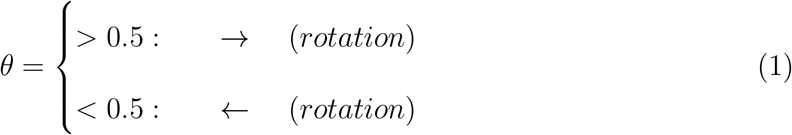

The currently perceived direction at timepoint *t* is defined by:

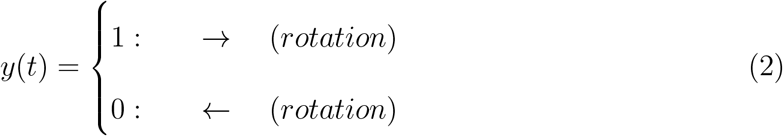

We manipulate the level of sensory information by changing the fraction of dots associated with a stereo-disparity signal. This is captured by a Gaussian distribution *Disambiguation* 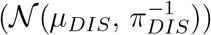. The direction of disambiguation at timepoint *t* is represented by *μ_Dis_*:

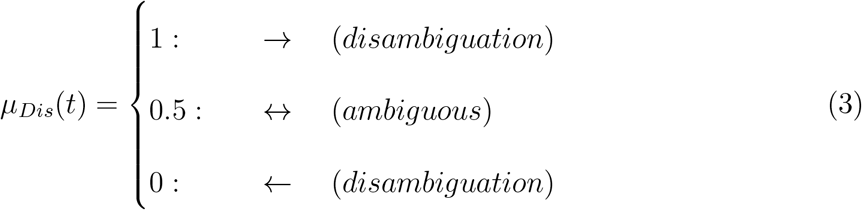

*π_DIS_* represents the participants’ sensitivity to disambiguating stimulus information. The amount of disambiguating stimulus information was varied systematically in 120 sec blocks of visual stimulation. The signal-to-ambiguity ratio (*SAR*) was defined by the fraction of stimulus dots that carried a 3D cue (level **D1**: 0.15, **D2**: 0.30, **D3**: 0.45, **D4**: 0.60, **D5**: 0.75 and **D6**; 1.00). If set to zero, *π_DIS_* is removed from the model.

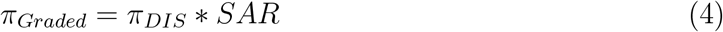

Furthermore, our model assumes that an implicit prior belief in the stability of the visual environment controls the frequency of changes in conscious experience during bistability. The mean of the Gaussian distribution “stability” 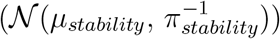 is determined by the perceptual state indicated by the participants at the overlap preceding timepoint *t*:

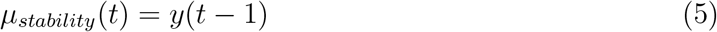

*π_stability_* describes the impact of the “stability” prior on perceptual state. If a change in conscious experience occurred at the preceding overlap (*t* = *t*_0_), *π_stability_* (*t*) is set to the initial stability precision *π_IPS_*:

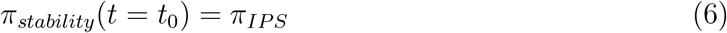

Inversion of our model during graded ambiguity allows for the estimation of *π_IPS_*. If fixed to zero, the parameter is removed from the model.

If no perceptual event occurred at the preceding overlap (*t* ≠ *t*_0_), we calculate *π_stability_* (*t*) by updating the previous precision of the stability prior *π_stability_*(*t* – 1) with a precision-weighted prediction error (PE). The precision of the prediction error (*π_ERROR_*) reflects how quickly *π_stability_* decays over time and is estimated as a free parameter:

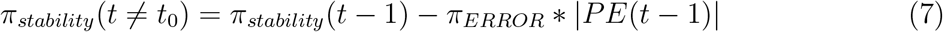

By combining the stability prior 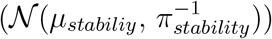 with the signal-to-ambiguity-adjusted likelihood 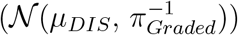, we adjust the density ratio *r* of the posterior *P*(*θ*) for the two peak locations *θ*_0_ = 0 and *θ*_1_ = 1:

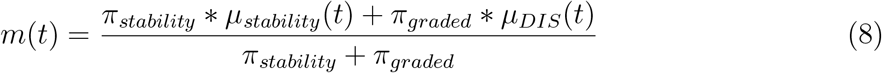

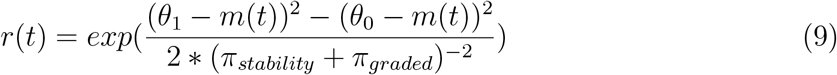

The posterior probability of right-ward rotation predicts the perceptual response *y*(*t*):

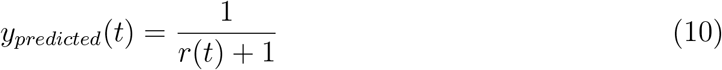

We infer on the free parameters (*π_DIS_, π_ERROR_, π_IPS_*) by optimizing the model with regard to the difference between the prediction and the actual perceptual response (*y_predicted_* and *y*). Once a new percept *y*(*t*) has been established, we compute the residual evidence for the alternative perceptual interpretation. This model quantity reflects a quantitative representation of dynamic changes in perceptual conflict. Given the inspiration of our model by predictive coding, we refer to this quantity as the *perceptual prediction error PE*(*t*):

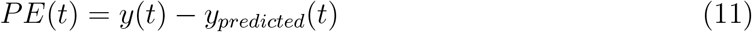

#### 9.2.2 Model inversion

For model inversion, we used a free energy minimization approach^66^, which maximised a lower bound on the log-model evidence for the individual participants’ data. We modeled *π_IPS_*, *π_ERROR_* and *π_DIS_* either as free parameters defined by log-normal distributions or fixed these entities to zero, thereby effectively removing them from the model. We optimised parameters using quasi-Newton Broyden-Fletcher-Goldfarb-Shanno minimisation as implemented in the HGF3.0 toolbox (TAPAS toolbox, http://www.translationalneuromodeling.org/hgf-toolbox-v3-0/).

For ambiguous visual stimulation, parameters were inverted using the following priors: *π_IPS_* = prior mean of log(2) and prior variance of 1; *π_ERROR_* = prior mean of log(1) and prior variance of 0.1. For graded ambiguity, prior means for *π_IPS_* and *π_ERROR_* were defined by the posterior estimates obtained from the preceding ambiguous runs. Prior variance was reduced to 0.01 for *π_IPS_* and to 0.001 for *π_ERROR_. π_DIS_* was estimated with a prior mean of log(2) and a prior variance of 1.

We used the inverted models for model-based fMRI in experiment E1 (posterior parameter estimates: *π_IPS_* = 2.83 ± 0.22; *π_ERROR_* = 0.7 ± 0.08) and E2 (*π_IPS_* = 2.25 ± 0.13; *π_ERROR_* = 0.57 ± 0.09; *π_DIS_* = 1.05 ± 0.15). Relative to model variants in which free parameters were systematically removed, models incorporating the full set of parameters (Ambiguity: *π_IPS_* and *π_ERROR_*, Graded ambiguity: *π_IPS_*, *π_ERROR_* and *π_DIS_*) were superior in explaining the participants’ behaviour (protected exceedance probability E1: 100%; E2: 99.98%). Furthermore, posterior model parameters were uncorrelated, indicating successful model inversion in E1 (*π_IPS_* to *π_ERROR_*: *ρ* = −0.18, *p* = 0.32) and E2 (*π_IPS_* to *π_ERROR_*: *ρ* = −0.35, *p* = 0.13; *π_IPS_* to *π_DIS_*: *p* = 0.17, *p* = %^47^; *π_ERROR_* to *π_DIS_*: *P* = 0.3, *P* = 0.2).

Based on previous work^14^, our model-based fMRI approach focused on perceptual prediction errors, since this model quantity provides a dynamic and quantitative representation of perceptual conflict.

#### 9.2.3 Simulation

To visualize the predictions of our model, we simulated experiment E2 (one run of ambiguous stimulation; three runs of graded ambiguity across six levels of sensory evidence D1 to D6) for a total of 100 hypothetical participants. Parameters for simulation were drawn randomly between the 30% and 70% quantile of posterior parameters estimated in the behavioral pretest (*π_IPS_* = 2.23 ± 0.14, *π_ERROR_* = 0.36 ± 0.07; *π_DIS_* = 1.73 ± 0.30).

As expected, the distribution of simulated phase durations (Supplementary Figure S10A) obtained during ambiguous stimulation showed a sharp rise and long tail. It was best fit by a gamma distribution (Bayesian Information Criterion = 3.83 × 10^4^; shape = 1.66, rate = 0.14) as compared to a lognormal (BIC = 3.85 × 10^4^) and a normal distribution (BIC = 4.11 × 10^4^).

When simulating graded ambiguity, we observed that disambiguating stimulus information biased the model predictions toward congruent perceptual states (Supplementary Figure S10B). This congruency effect was stronger at higher levels of signal-to-ambiguity (F(495) = 195.1, p = 1.49 × 10^-114^, *BF*_10_ = 4.18 × 10^120^). Simulated prediction errors signaled elevated perceptual conflict during incongruent as opposed to congruent perceptual states (main effect of *Congruency*: F(1.09 × 10^3^) = 4.15 × 10^3^, p = 0, *BF*_10_ = 7.16 × 10^275^, Figure S10C). Differences in prediction errors between congruent and incongruent perceptual states scaled with the signal-to-ambiguity ratio (interaction between *Congruency* and *Signal-to-Ambiguity*: F(1.09 × 10^3^) = 148.71, p = 2.1 × 10^-120^, *BF*_10_ = 1.99 × 10^116^). We also observed a main effect of *Signal-to-Ambiguity* (F(1.09 × 10^3^) = 81.43, p = 1.07 × 10^-72^, *BF*_10_ = 1.28 × 10^42^).

Thus, when simulating from this computational model, we observed that the model’s predictions closely followed the behavioral characteristics of both full and graded ambiguity.

### 9.3 Decoding the contents of conscious experience

We used multi-variate pattern analysis to isolate brain regions that contained information about the dynamic *contents of conscious experience*. This was achieved in a standard cross-validated leave-one-run-out searchlight decoding analysis^67^. This approach classified between two regressors that represented the alternative stimulus interpretations, i.e., leftward and rightward rotation. Thereby we predicted, based on brain activity, which of the two perceptual states was dominant or suppressed at a given point in time. In both experiments E1 and E2, searchlight analysis was carried out with a radius of 8 voxels (unsmoothed data), using a support vector machine^68^ as a binary classifier.

In both experiments, information about the current content of conscious experience was primarily represented in visual and parietal brain region (Supplementary Figure S8A). To test whether the representation of perceptual conflict in IFC and V5/hMT+ overlapped with brain regions that contained information about the current content of conscious experience, we computed average classification accuracy in voxels within V5/hMT+ and IFC (see section 7.3.4 for ROI-definition).

Classification accuracy was significantly above chance in V5/hMT+ for data from both E1 (T(31) = 2.35, p = 0.03, *BF*_10_ = 2.03) and E2 (T(19) = 3.02, p = 6.97 × 10^-3^, *BF*_10_ = 6.93, Supplementary Figure S8B). In IFC, we found no conclusive evidence for above-chance classification in E1 (T(31) = 1.95, p = 0.06, *BF*_10_ = 1) or E2 (T(19) = 1.92, p = 0.07, *BF*_10_ = 1.08). Hence, reliable decoding of the contents of conscious experience was possible for V5/hMT+, but not IFC.

### 9.4 Perceptual uncertainty and temporal balances during graded ambiguity

Our imaging analysis of E2 strongly suggested that IFC detected dynamic changes in perceptual conflict during graded ambiguity. In addition to model-based fMRI, this interpretation was supported by IFC reflecting both a main effect of *Congruency* and an interaction between *Congruency* and *Signal-to-Ambiguity*. Here, we provide additional control analyses to rule out that differences in perceptual uncertainty and temporal imbalance of perceptual states at different levels of *Signal-to-Ambiguity* may have confounded these fMRI results.

Firstly, since previous research has associated IFC with unclear percepts during bistability^15^, we asked whether differences in perceptual uncertainty between congruent and incongruent perceptual states could provide an alternative interpretation of our findings. In experiment E2, unclear perceptual states were extremely rare (0.29 ± 0.13%) and unaffected by levels of *Signal-to-Ambiguity* (F(95) = 1.4, p = 0.23), indicating that participants were able to discriminate between the alternative stimulus interpretations at all times. In an additional control experiment (Run 3 of E2, Supplementary Figure S1B), we obtained ratings of perceptual uncertainty associated with congruent and incongruent perceptual states (Supplementary Figure S11). Perceptual uncertainty was low for both congruency (average rating: 1.2 ± 0.05, range: 1 to 4) and incongruency (1.4 ± 0.06) to disambiguating stimulus information. Incongruent perceptual states were associated with an increase in perceptual uncertainty (Chi-square = 19.43, p = 3.5 × 10^-3^, *BF*_10_ = 27.3). Importantly, however, such differences in perceptual uncertainty were unaffected by levels of *Signal-to-Ambiguity* (Chi-square = 7.41, p = 0.19, *BF*_10_ = 5.53 × 10^-5^).

Secondly, we asked whether the temporal imbalance between congruent and incongruent perceptual states might have confounded our imaging results. In a fMRI control experiment (Run 8 of E2, Supplementary Figure S1B), participants reported the direction of motion (left vs. right) of unambiguous 2D dot fields. Across six levels of *Temporal Imbalance*, we matched the *Relative Duration* of perceptual outcomes to the effects of signal-to-ambiguity on perceptual congruency. Even at a lenient threshold of p < 0.001 (uncorrected voxel-wise whole-brain analysis), we did not observe a main effect of *Relative Duration* or an interaction between *Relative Duration* and *Temporal Imbalance* in any of the regions implicated in the representation of perceptual conflict.

In sum, neither differences in perceptual uncertainty nor temporal imbalances between congruent and incongruent perceptual states provided alternative explanations for the modulation of neural activity in V5/hMT+ and IFC by perceptual conflict.

### 9.5 Effects of TMS on the stability, the uncertainty and the report of conscious experience during bistable perception

In our analysis of E3, we have focused on differences in phase duration, unclear perceptual states and RTs associated with TMS to either IFC or vertex. We observed that TMS-induced virtual lesions in IFC were associated with a pronounced prolongation of phase durations. Importantly, we found no evidence for an influence of IFC on either the uncertainty or the report of perceptual states.

Here, we confirmed these results in linear mixed effects modeling. To this end, we analyzed the counter-balanced TMS experiment with respect to two factors: *Session* (IFC vs. vertex) and *Stimulation* (post vs. pre). We used linear mixed effects modeling (lmer R-package) with fixed effects for *Session* and *Stimulation*. Random effects were defined for individual participants.

The relative prolongation of phase durations after TMS to IFC was reflected by a significant interaction between the factors *Stimulation* and *Session* (F(87) = 6.35, p = 0.01, *BF*_10_ = 3.8). We did not observe a main effect of *Stimulation* (F(87) = 2.47, p = 0.12, *BF*_10_ = 0.54) or *Session* (F(87) = 0.11, p = 0.29, *BF*_10_ = 0.2; Supplementary Figure S1D).

With respect to unclear perceptual states (Supplementary Figure S12), we found no main effect of *Session* (F(58) = 0.34, p = 0.56, *BF*_10_ = 0.41) or *Stimulation* (F(58) = 0.35, p = 0.56, *BF*_10_ = 0.21) and no between-factor interaction (F(58) = 1.12, p = 0.3, *BF*_10_ = 0.35). Likewise, when assessing a potential modulation of RTs by TMS (Supplementary Figure S3D), we found no main effect *Session* (F(58) = 0.07, p = 0.8, *BF*_10_ = 0.21) nor *Stimulation* (F(58) = 0.07, p = 0.79, *BF*_10_ = 0.21) nor a between-factor interaction (F(58) = 0.52, p = 0.47, *BF*_10_ = 0.35).

Linear mixed effects modeling thus corroborated that TMS-induced virtual lesions in IFC had a selective effect on the occurrence of changes in conscious experience during bistable perception.

### 9.6 Effects of stimulus exposure on the dynamics of conscious experience

In binocular rivalry, changes in conscious experience evoked by binocular conflict have been shown to occur more frequently when participants were exposed to the sensory ambiguity repeatedly over several days^69^. Likewise, in E1 and E3, we observed a shortening of phase durations elicited by structure-from-motion with increasing stimulus exposure (F(321.65) = 5.9, p = 9.71 × 10^-9^, *BF*_10_ = 1.18 × 10^6^; see Supplementary Figure S2C).

To make sure that the effects attributed to TMS-induced virtual lesions were not confounded by this effect of exposure, we re-analyzed the behavioral data from the counter-balanced TMS-experiment with respect to three factors: *Exposure* (increasing from Run 1 to Run 8), *Session* (IFC vs. vertex) and *Stimulation* (post vs. pre).

Linear mixed effects modeling confirmed the main effect of stimulus exposure on phase duration (F(202.03) = 14.43, p = 1.93 × 10^-4^) in the course of the TMS-experiment. As a reflection of the effect of TMS on the stability of conscious experience, we found a highly significant interaction between the factors *Session* (IFC vs. vertex) and *Stimulation* (post vs. pre; F(202.03) = 11.53, p = 8.25 × 10^-4^), a main effect of *Stimulation* (F(202.03) = 4.85, p = 0.03), but no main effect of *Session* (F(202.03) = 0.12, p = 0.73). However, the effects of *Exposure* were independent of the effects of *Stimulation* and *Session* (*BF*_10_ = 23.76). Linear mixed effects modeling did not reveal a two-way interaction between the factors *Exposure* and *Session* (F(202.03) = 0.03, p = 0.87) or *Stimulation* (F(28.01) = 3.73 × 10^-3^, p = 0.95) and no three-way interaction (F(202.03) = 0.28, p = 0.59).

We thus did not observe any evidence for a confounding of TMS-related effects by effects introduced by stimulus exposure.

### 9.7 Regression toward the mean

As a consequence of the independence between the long-term speeding induced by stimulus exposure and the IFC-TMS-induced stabilization of conscious experience, pre-stimulation phase duration differed between vertex- and IFC-in runs E3(5) (IFC: 19.82 ± 3.69 sec; vertex: 25.22 ± 4.65 sec; one-sided two-sample t-test: T(26.63) = −0.91, p = 0.19) and E3(6) (IFC: 19.38 ± 4.83 sec; vertex: 24.09 ± 4.06 sec; T(26.52) = −0.75, p = 0.23). One may thus assume that, by virtue of *regression toward the mean* (RTM), such differences at baseline might have confounded our analyses of TMS-related effects.

However, due to the high correlation between pre- and post-TMS-measurements, variance attributable to RTM (i.e., 100 * (1 – *ρ_Spearman_*)%) was low for TMS to both vertex (13.26%) and IFC (11.43%). As an additional control analysis, we applied Mee and Chua’s test^70^ to differentiate between the RTM effect and additional TMS effects *τ*. For TMS to IFC, we found significant TMS effects over and above RTM estimated *τ* = 7.98 sec; T = 4.83, p = 4.38 × 10^-5^). As expected, control stimulation to the vertex was not associated with any significant additional effects on phase duration (estimated *τ* = −1.53 sec; T = −0.73, p = 0.47). In sum, these analyses provided strong evidence against a potential confound of the observed TMS effects by RTM.

### 9.8 Brain-behavior associations

Support vector regression (SVR) applied in a whole-brain searchlight approach revealed that localized patterns of BOLD activity in IFC predicted the individual effect virtual IFC-lesions on conscious experience (Figure 4E, upper panel). In a first control analysis, we asked whether this brain-behavior association was specific to virtual lesions in IFC. To this end, we used SVR to predict inter-individual differences in the effect of control stimulation at vertex based on the neural correlates of perceptual prediction errors. This analysis did not reveal a significant assocation between patterns of BOLD-activity in IFC and individual post-pre differences in phase duration after control stimulation at vertex (*p_FWE_* < 0.05).

In a second set of control analyses, we asked whether inter-individual differences in phase duration prior to IFC stimulation may represent a potential confound for the observed association between IFC activity and post-pre differences in phase duration induced by virtual IFC-lesions. On the level of behavior, we found that participants with longer pre-stimulation phase duration at baseline showed a larger post-pre difference in phase duration after stimulation at IFC (*ρ* = 0.44, *p* = 0.02), but not after control stimulation at vertex (*ρ* = –0.1, *p* = 0.6). This argued against the possibility that differences in pre-stimulation baseline may have affected post-pre differences in phase duration irrespective of whether IFC activity was disrupted by TMS.

On the neural level, we tested whether localized patterns of BOLD-activity in IFC were predictive of inter-individual differences in phase duration prior to stimulation of IFC. SVR did not reveal any significant association between the neural representation of perceptual prediction errors in IFC and pre-stimulation phase duration (*p_FWE_* < 0.05).

In sum, these analyses argue against the view that the observed association between the neural representation of perceptual conflict in IFC and individual effects of virtual IFC-lesions on conscious experience was confounded by differences in phase duration at baseline.

### 9.9 Supplementary Figures

**Supplementary Figure S1.**
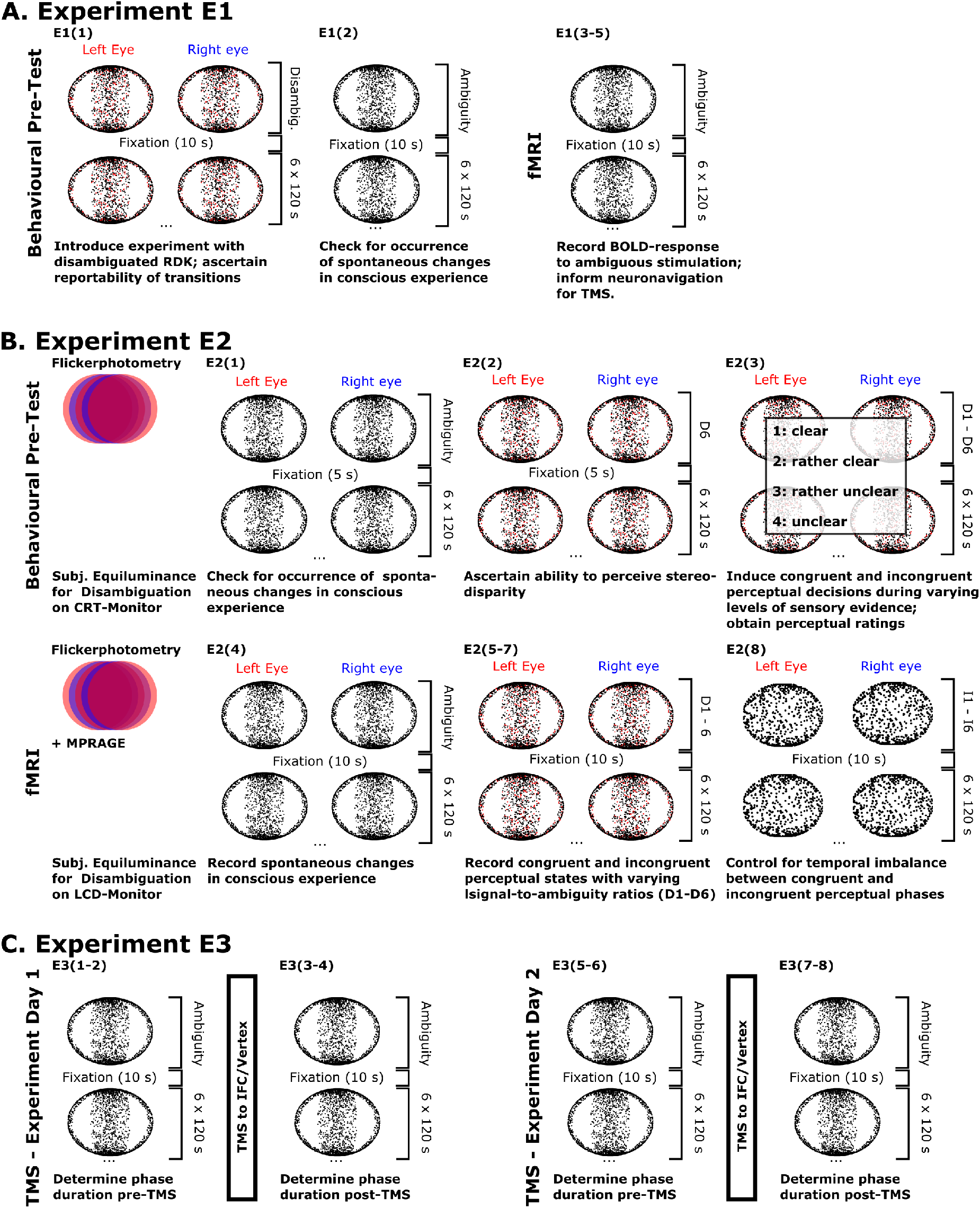
Experimental Paradigm. **A. Experiment E1.** In E1, we investigated how IFC responded to the perceptual conflict inherent in ambiguous visual stimulation. The experimental protocol started with a behavioral pre-test, which familiarized the participants with the experiment. **Run 1** tested adequate performance in reporting changes in direction of rotation of a fully disambiguated RDK (probability: 15% per overlapping configuration). **Run 2** checked whether participants perceived spontaneous changes in conscious experience during ambiguous visual stimulation. In a separate experimental session, we collected fMRI-data during three runs of ambiguous structure-from-motion (**Runs 3-5**). This allowed us to assess how IFC responded to the perceptual conflict during bistable perception. In addition, the fMRI data were used to localize change-related BOLD activity in IFC for neuronavigated inhibitory TMS (see E3). **B. Experiment E2.** In E2, we tested how neural activity in IFC responded to varying levels of sensory ambiguity. In a behavioral pretest, we tested whether participants perceived a sufficient number of changes in conscious experience during full ambiguity (**Run 1**) and assessed their ability to perceive 3D cues during full disambiguation (**Run 2**). **Run 3** introduced graded ambiguity. Here, a fraction of the dots composing the RDK (highlighted in red) were shifted between the two monocular channels. The conditions D1 to D6 were defined by the percentage of disambiguated dots (signal-to-ambiguity ratio, 15%, 30%, 45%, 60%, 75% and 100%) and occurred in random order. We determined the frequency of exogenous stimulus changes during graded ambiguity based on the frequency of changes in conscious experience during full ambiguity. In Run 3, we tested the effects of signal-to-ambiguity on the proportion of perceptual states congruent to the disambiguating stimulus information. Moreover, we obtained ratings of perceptual uncertainty for both congruent and incongruent perceptual states across six levels of signal-to-ambiguity. To test for effects of signal-to-ambiguity on neural activity in IFC, we measured BOLD activity during one run of full ambiguity (**Run 4**) and three runs of graded ambiguity (**Runs 5-7**). In **Run 8**, we conducted a final control experiment assessing the effects of temporal imbalance introduced between congruent and incongruent perceptual states across conditions D1-D6. Here, we presented leftward and rightward planar dot motion in six conditions of increasing temporal imbalance (I1-I6). All experiments in E2 used filter-glasses to present 3D stimuli. We adjusted the subjective luminance of the red (left eye) and blue (right eye) channel using heterochromatic flicker photometry. **C. Experiment E3.** E1 and E2 established that neural activity in right-hemispheric IFC signals the accumulating conflict between conscious experience and the underlying sensory information. In E3, we probed a causal contribution of IFC activity to conscious experience. We re-invited the participants of E1 to two additional TMS-session scheduled on consecutive days (Day 1: **Runs 1-4**; Day 2: **Runs 5-8**). Using online neuronavigation based on the fMRI experiment in E1, we created virtual lesions in IFC and a control region at the cranial vertex (order counter-balanced across participants). We assessed the effects of virtual lesions on the dynamics and content of conscious experience by comparing pre-with post-stimulation runs.

**Supplementary Figure S2.**
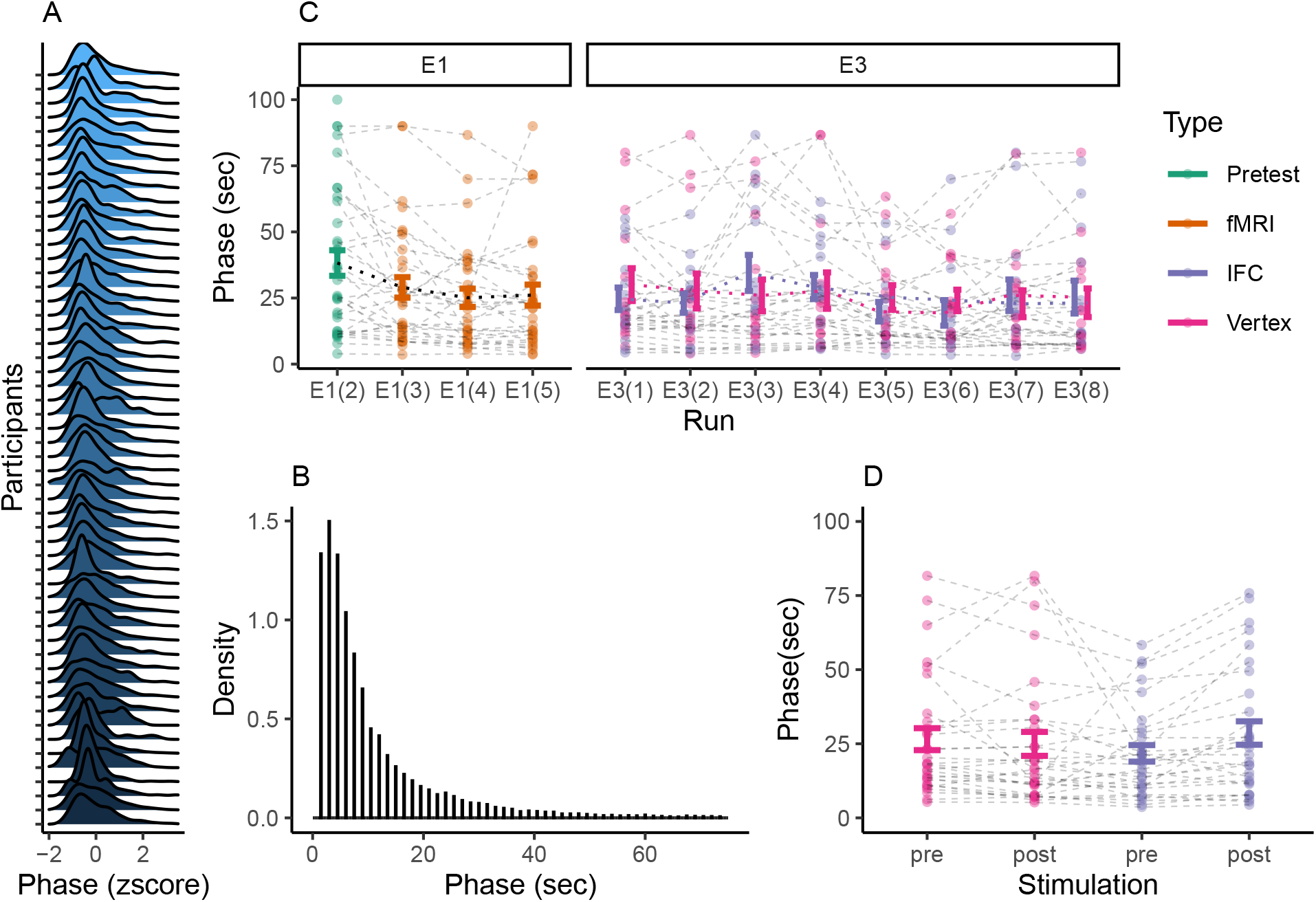
Phase Duration. **A. Individual distributions of phase duration.** In individual participants (E1-E3), the distribution of phase durations (z-scored for visualization) followed the sharp rise and long tail that is typically observed in perceptual bistability. **B. Aggregated phase durations.** When aggregated across participants, the distribution of phase durations (multiples of the 1.5 sec inter-overlap interval) was again characterized by a sharp rise and long tail. This pattern was best fit by a log-normal distribution (Bayesian Information Criterion = 1.79 × 10^5^; log(mean) = 1.91, log(sd) = 0.95) as compared to a gamma (BIC = 1.85 × 10^5^) and a normal distribution (BIC = 2.18 × 10^5^). **C. Effects of stimulus exposure on phase duration.** Across the experimental runs of E1 and E3, we observed a significant decrease in phase duration with increasing exposure to the bistable stimulus (F(321.65) = 5.9, p = 9.71 × 10^-9^, *BF*_10_ = 1.18 × 10^6^; see dotted lines). When assessing the evolution of phase duration across the counter-balanced TMS-experiment (E3; blue dotted line: IFC-stimulation first; red dotted line: vertex-stimulation first), linear mixed effects modeling again showed a main effect of *Exposure* (F(202.03) = 14.43, p = 1.93 × 10^-4^). As expected, we also found a highly significant interaction between the factors *Session* (IFC vs. vertex) and *Stimulation* (post vs. pre; F(202.03) = 11.53, p = 8.25 × 10^-4^), a main effect of *Stimulation* (F(202.03) = 4.85, p = 0.03), but no main effect of *Session* (F(202.03) = 0.12, p = 0.73). Importantly, we did not observe any two-way interactions between the factors *Exposure* and *Session* (F(202.03) = 0.03, p = 0.87) or *Stimulation* (F(28.01) = 3.73 × 10^-3^, p = 0.95) and no three-way interaction (F(202.03) = 0.28, p = 0.59). This argued for the independence of the effects of exposure from TMS-related changes in conscious experience (*BF*_10_ = 23.76). **D. Effects of TMS on phase duration.** Here, we display phase duration collapsed across the two factors *Stimulation* (post vs. pre) and *Session* (IFC vs. vertex) in E3. Phase durations showed no main effect of *Stimulation* (F(87) = 2.47, p = 0.12, *BF*_10_ = 0.54) and no main effect of *Session* (F(87) = 0.11, p = 0.74, *BF*_10_ = 0.2). As expected, we found a highly significant between-factor interaction (F(87) = 6.35, p = 0.01, *BF*_10_ = 3.8), reflecting the relative prolongation of phase duration after TMS to IFC (see Figure 4A
).

**Supplementary Figure S3.**
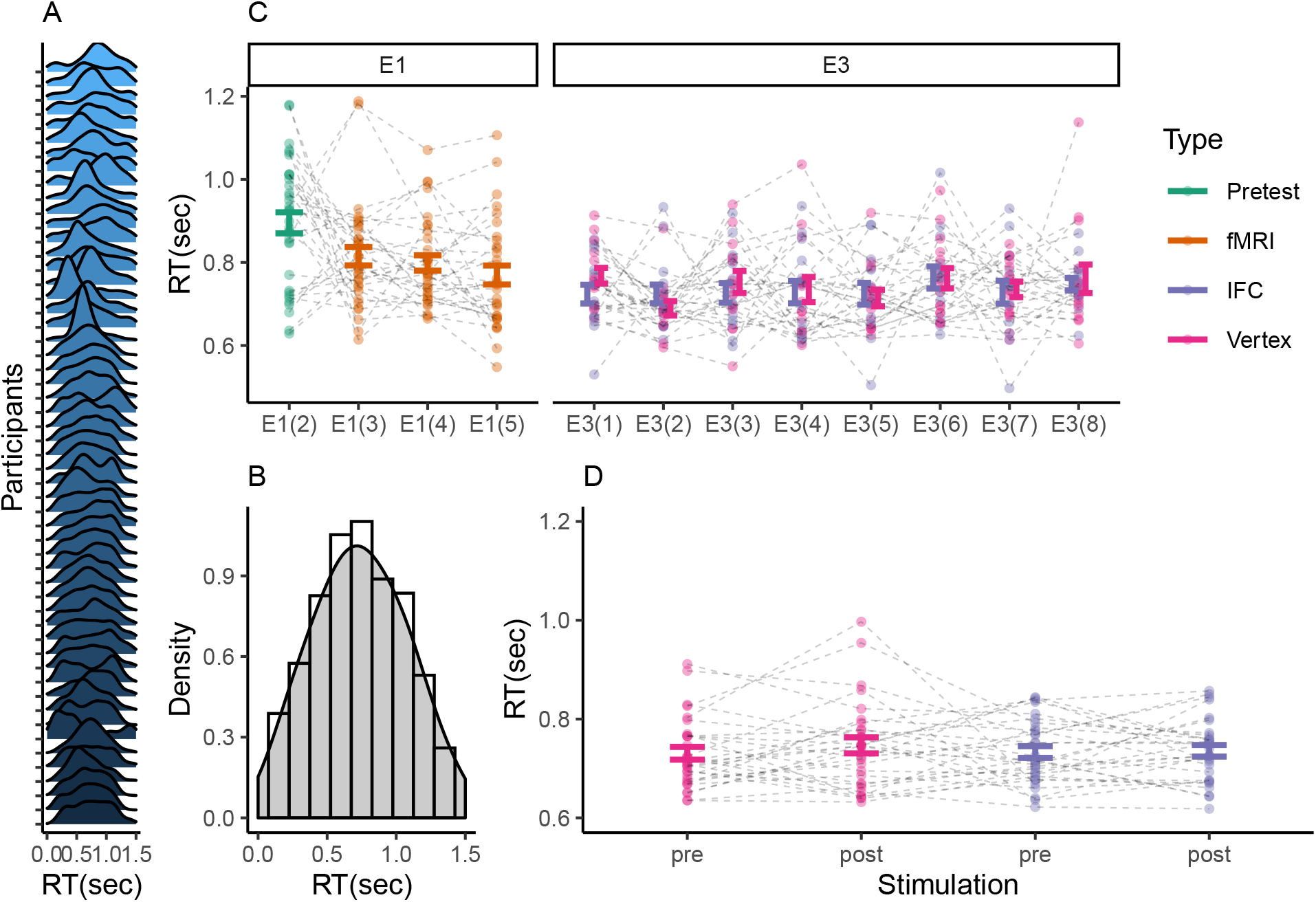
Response Times. **A. Discontinuous bistable perception.** Here, we depict the distribution of RTs for the report of changes in conscious experience in individual participants across E1, E2 and E3. Timepoints are displayed relative to the onset of the preceding overlap. Response times were distributed non-uniformly across the inter-overlap interval (chi-square test against uniformity: p < 0.05 in 55 of 55 participants). This ensured that changes in conscious experience were indeed confined to overlapping stimulus configurations, indicating that conscious experience unfolded in a discrete sequence of *perceptual states*. **B. Collapsed RTs.** Across participants, aggregated RTs (0.81 ± 0.05 sec) were best fit by a normal distribution (BIC = 1.77 × 10^4^, mean = 0.74, sd = 0.33) as compared to a gamma (BIC = 2.1 × 10^4^) or a lognormal distribution (BIC = 2.8 × 10^4^). **C. Effects of stimulus exposure on RTs.** Across the experimental runs of E1 and E3, prolongated exposure to structure-from-motion was associated with a reduction in response times (F(328.25) = 9.29, p = 1.55 × 10^-14^). **D. Effects of TMS on RTs.** In E3, RTs did not vary according to the factors *Stimulation* (post vs. pre; F(58) = 0.99, p = 0.32, *BF*_10_ = 0.27) and *Session* (IFC vs. vertex; F(58) = 0.07, p = 0.8, *BF*_10_ = 0.21). Importantly, we did not observe a significant between-factor interaction (F(58) = 0.52, p = 0.47, *BF*_10_ = 0.3, see also Figure 4C).

**Supplementary Figure S4.**
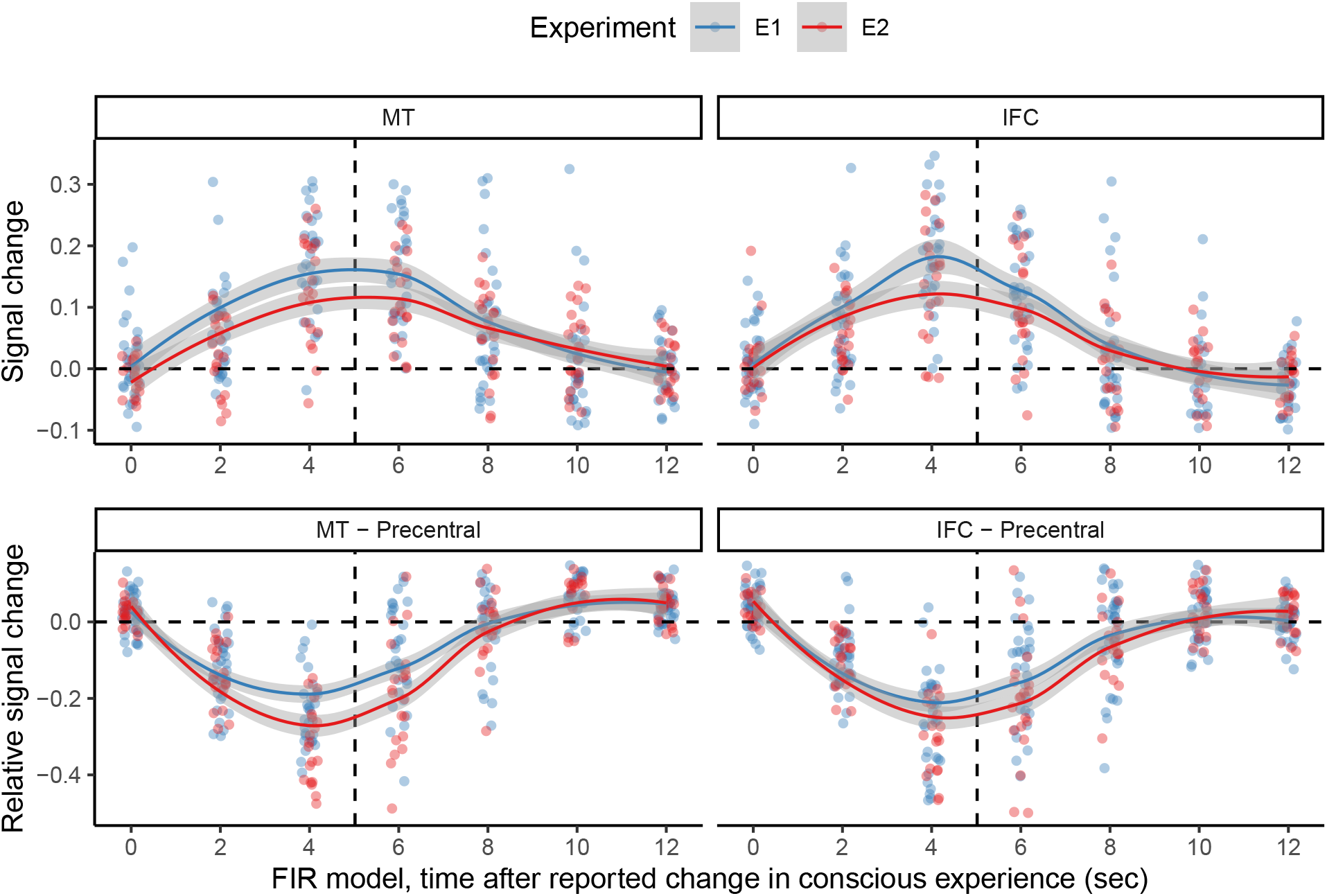
Conflict-related BOLD signals around the time of changes in conscious experience. In both E1 (blue) and E2 (red), neural activity in MT and IFC (upper panel) gradually increased towards the peak of change-related activity (vertical dotted line). Relative to a motor-related control region in left precentral gyrus (see Table S1), BOLD signals (lower panel) were significantly elevated *prior* to the surge of change-related activity (timepoint 0, i.e., timing of the overlap preceding a reported change in conscious experience) in both MT (E1: T(32) = 2.42, p = 0.01, *BF*_10_ = 2.29; E2: T(19) = 3.38, p = 1.58 × 10^-3^, *BF*_10_ = 13.59; two-sided t-test) and IFC (E1: T(32) = 4.25, p = 8.61 × 10^-5^, *BF*_10_ = 155.63; E2: T(19) = 3.55, p = 1.07 × 10^-3^, *BF*_10_ = 18.99). This result aligns with the hypthothesis that MT and IFC signal accumulating perceptual conflicts that lead up to changes in conscious experience. As expected, change-related activiy itsself was more pronounced in left precentral gyrus (timepoints 2, 4 and 6; all p < 0.001).

**Supplementary Figure S5.**
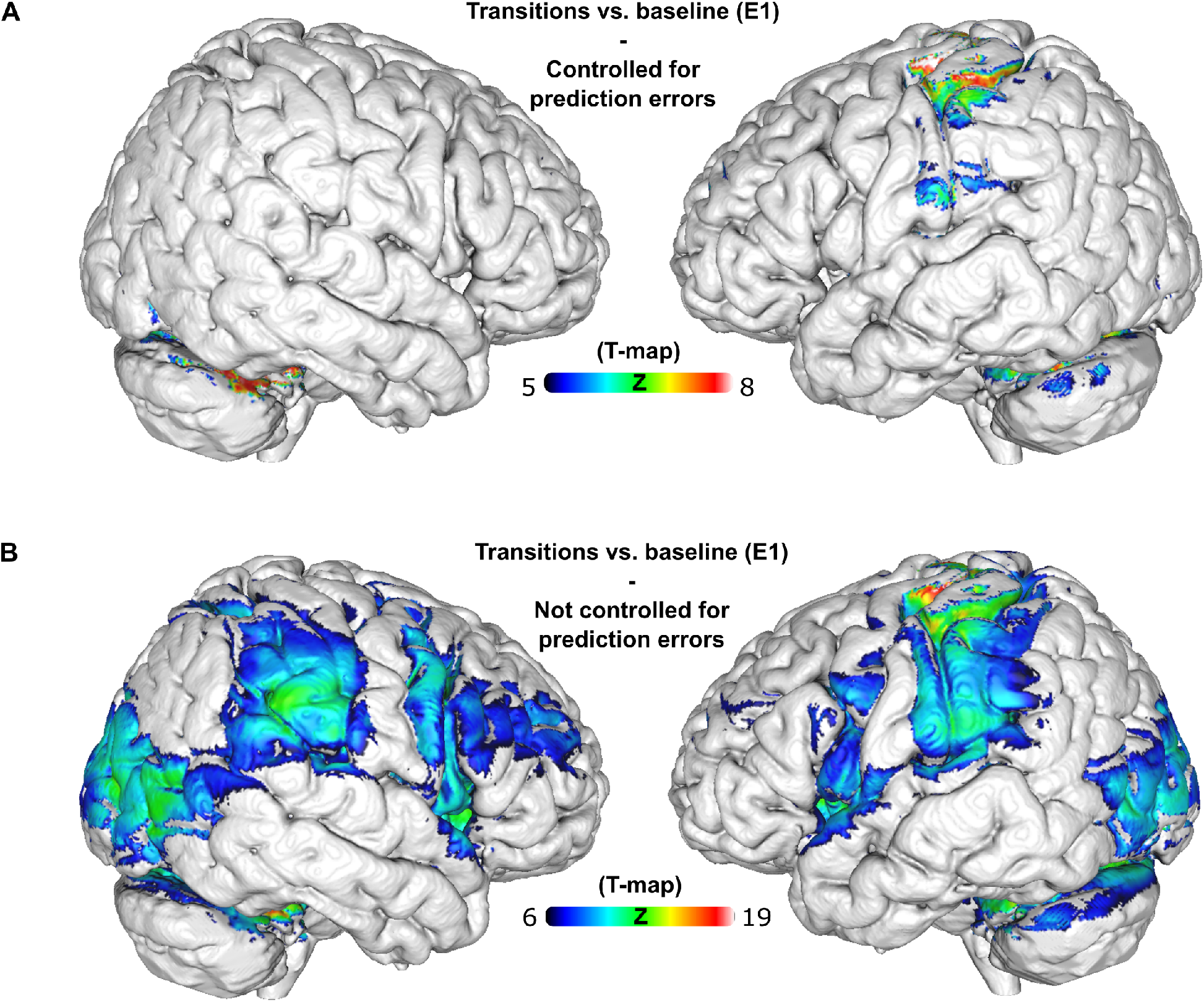
The neural correlates of reported changes in perception. **A. Controlled for perceptual prediction errors.** When analyzing the neural correlates of perceptual events while controlling for BOLD activity related to gradually accumulating perceptual prediction errors (GLM-PC), we found activations in bilateral cerebellum, left pre- and postcentral gyrus, bilateral midcingulate cortex and putamen, left insula, left IPL as well as left medial frontal gyrus (*p_FWE_* = 0.05). No significant clusters were observed in right-hemispheric IFC or V5/hMT+. **B. Not controlled for perceptual prediction errors.** When assessing the neural correlates of perceptual events without controlling for perceptual prediction errors (i.e., by deleting the prediction-error regressor from GLM-PC), we observed highly significant change-related activity in bilateral insula, right inferior frontal gyrus, bilateral V5/hMT+, bilateral cerebellum, left pre- and postcentral gyrus, bilateral midcingulate cortex, bilateral inferior parietal lobulus and left middle frontal gyrus (*p_FWE_* = 10^-6^). Hence, when studied in isolation of perceptual prediction errors, perceptual events did activate regions in right IFC (see Supplementary Figure S6 for a comparison of the explanatory power of change-related regressors and accumulating perceptual prediction errors with respect to BOLD activity).

**Supplementary Figure S6.**
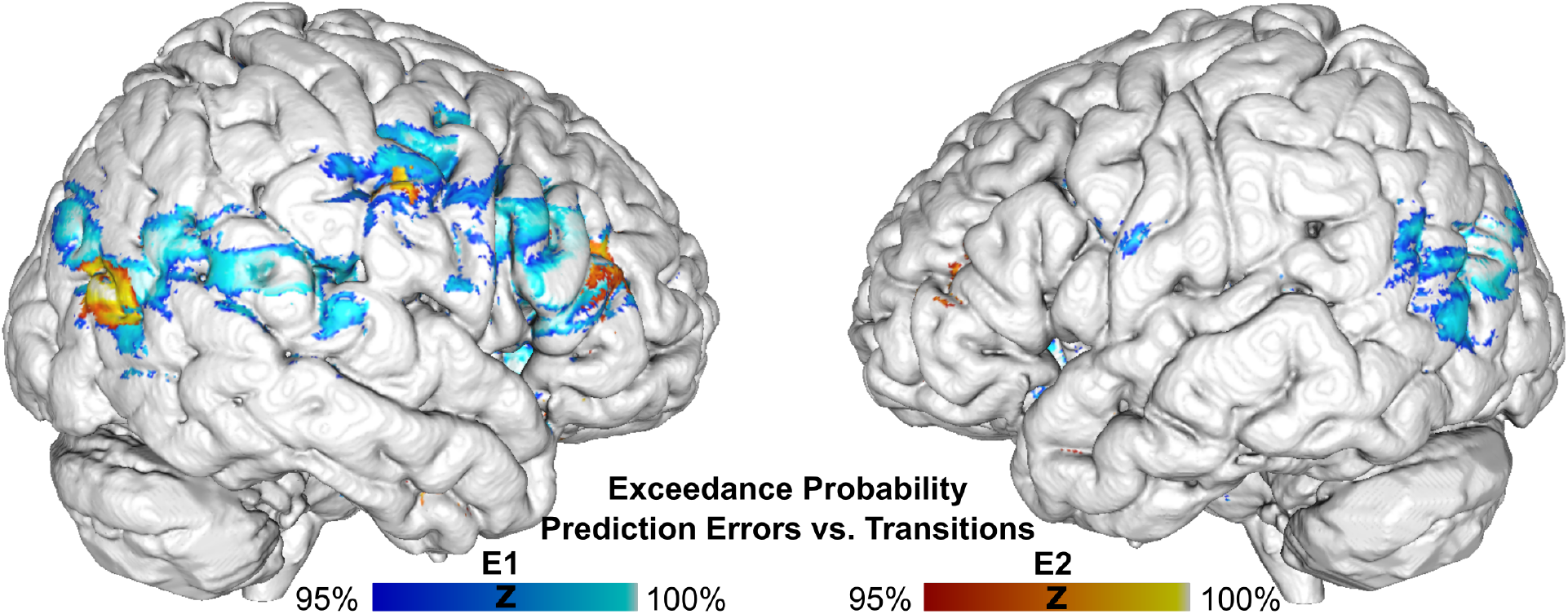
Posterior probability maps. We applied a Bayesian Posterior Probability Maps approach to compare the explanatory power of gradually accumulating perceptual prediction errors against the explanatory power of event-related regressors aligned with perceptual changes. Here, we display voxels where BOLD activity was better explained by gradually accumulating prediction errors at an exceedance probability above 95% (E1: blue heatmap; E2: red heatmap). Across both experiments, the posterior probability maps yielded converging evidence that neural signals from IFC and V5/hMT+ (as well as from additional parietal brain regions) were better explained by prediction-error related activity as compared to change-related activity.

**Supplementary Figure S7.**
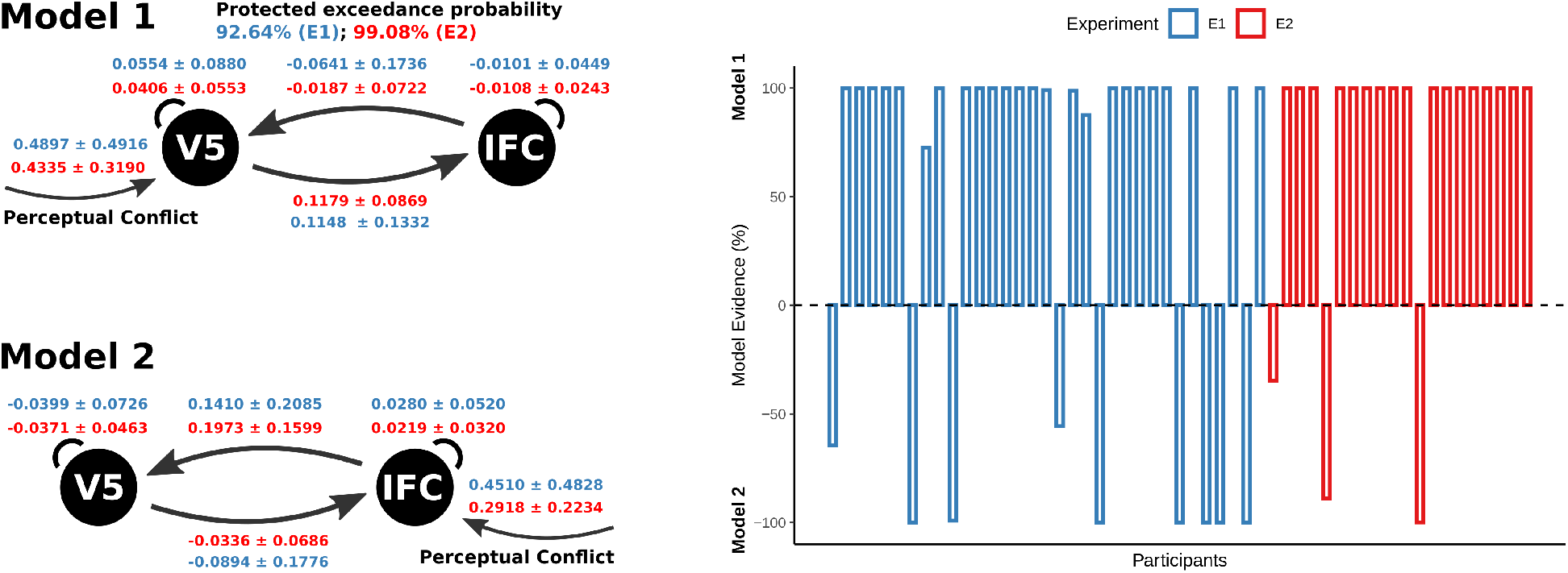
Dynamic causal modeling of perceptual conflict. Here, we used dynamic causal modelling^25^ to test whether the neural correlates of perceptual conflict were more likely to originate from V5/hMT+ or, alternatively, from IFC. To this end, we extracted eigenvariate timecourses from individual V5/hMT+- and IFC-ROIs (see Methods section 7.3.4) and constructed two linear, one-state-per-region, non-stochastic dynamic causal models (DCMs; see left panel for model layouts). Both Model 1 and Model 2 incorporated mutual connections between V5/hMT+- and IFC (A-matrix). In Model 1, perceptual conflict acted as a driving input to V5/hMT+ (C-matrix). By contrast, in Model 2, perceptual conflict was defined to drive activity in IFC. In each experiment, we estimated the two models for each session and participant and conducted group-level inference using random-effects Bayesian model comparison^71^. Across both experiments, we found that Model 1 clearly outperformed Model 2 (E1: 92.64% protected exceedance probability for Model 1; E2: 99.08%; see right panel for posterior model evidence in individual participants). Bayesian parameter averages (E1 in blue, E2 in red) were highly consistent across experiments, indicating that perceptual conflict acted as a positive driving factor into V5/hMT+, which, in turn, drove activity in IFC via feedforward effecive connectivity. Thus, dynamic causal modeling of both experiments provided converging evidence that perceptual conflict was most likely to originate from neural activity within V5/hMT+ and to reach prefrontal cortex via feedforward processing from V5/hMT+ to IFC.

**Supplementary Figure S8.**
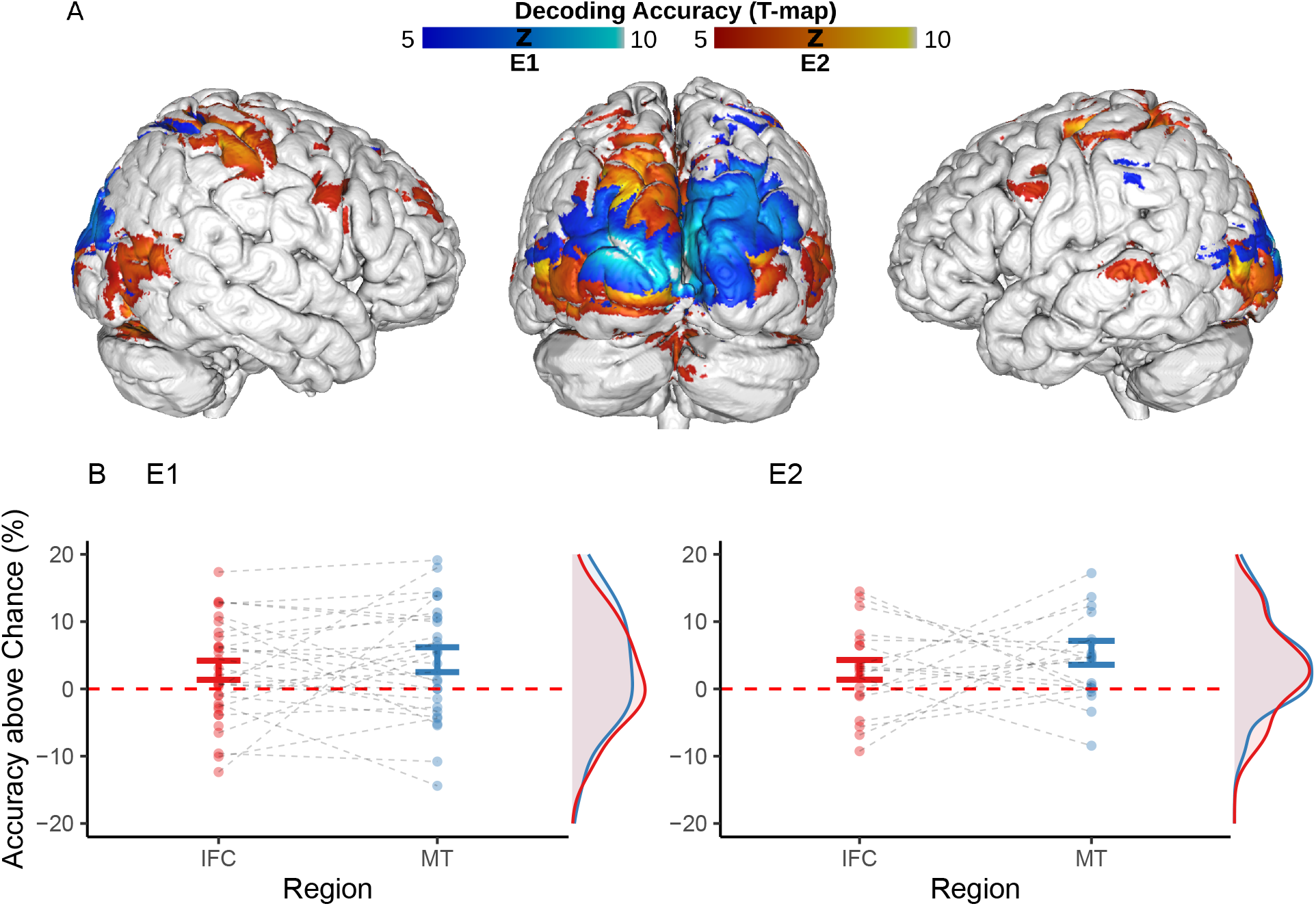
Decoding the contents of conscious experience. **A. Whole-brain searchlight.** Multi-variate pattern analysis of BOLD-data suggested that information about the current content of conscious experience was pre-dominantly represented in posterior brain regions such as visual cortex (including V5/hMT+) and parietal cortex (SPL and IPL). **B. Decoding accuracy in hMT and IFC.** In E1 (left panel), classification accuracy was significantly above chance in V5/hMT+ (T(31) = 2.35, p = 0.03, *BF*_10_ = 2.03). For IFC, we did not find conclusive evidence for above-chance classification (T(31) = 1.95, p = 0.06, *BF*_10_ = 1). Likewise, in E2 (right panel), we were able to decode the current contents of conscious experience from V5/hMT+ (T(19) = 3.02, p = 6.97 × 10^-3^, *BF*_10_ = 6.93). E2 yielded no conclusive evidence for above-chance classification in IFC (T(19) = 1.92, p = 0.07, *BF*_10_ = 1.08).

**Supplementary Figure S9.**
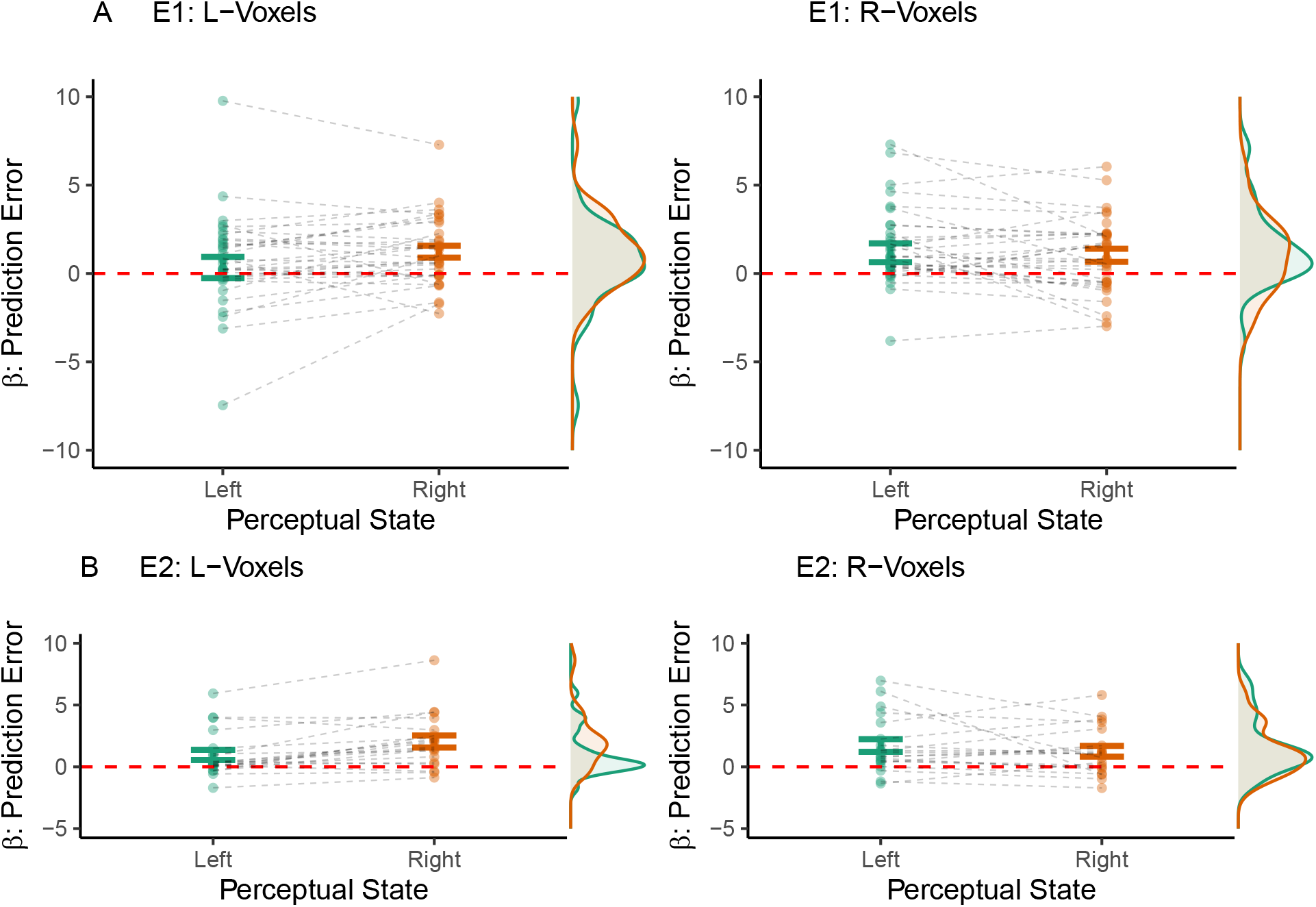
Perceptual prediction errors in voxels with reduced biases. **A. Experiment E1.** As shown above, voxels with biases toward one perceptual interpretation (T > 1) represented perceptual conflict more strongly when the corresponding perceptual interpretation was suppressed. Here, we show that this finding depended on the *strength* of voxel biases. To this end, we selected voxels with reduced biases toward left- or rightward illusory rotation (0 < T < 0.5). In these voxel populations, we re-tested the correlation of BOLD activity with perceptual prediction errors. In this analysis, the representation of perceptual states was not dissociated from the representation of perceptual conflict (L-voxels: T(32) = −1.83, p = 0.08, *BF*_10_ = 0.83; R-voxels: T(32) = 0.27, p = 0.79, *BF*_10_ = 0.19). **B. Experiment E2.** Again, we found no consistent evidence for a dissociation between the representation of perceptual states and representation of perceptual conflict in voxels with reduced biases (L-voxels: T(19) = −4.04, p = 7.01 × 10^-4^, *BF*_10_ = 49.88; R-voxels: T(19) = 0.98, p = 0.34, *BF*_10_ = 0.36).

**Supplementary Figure S10.**
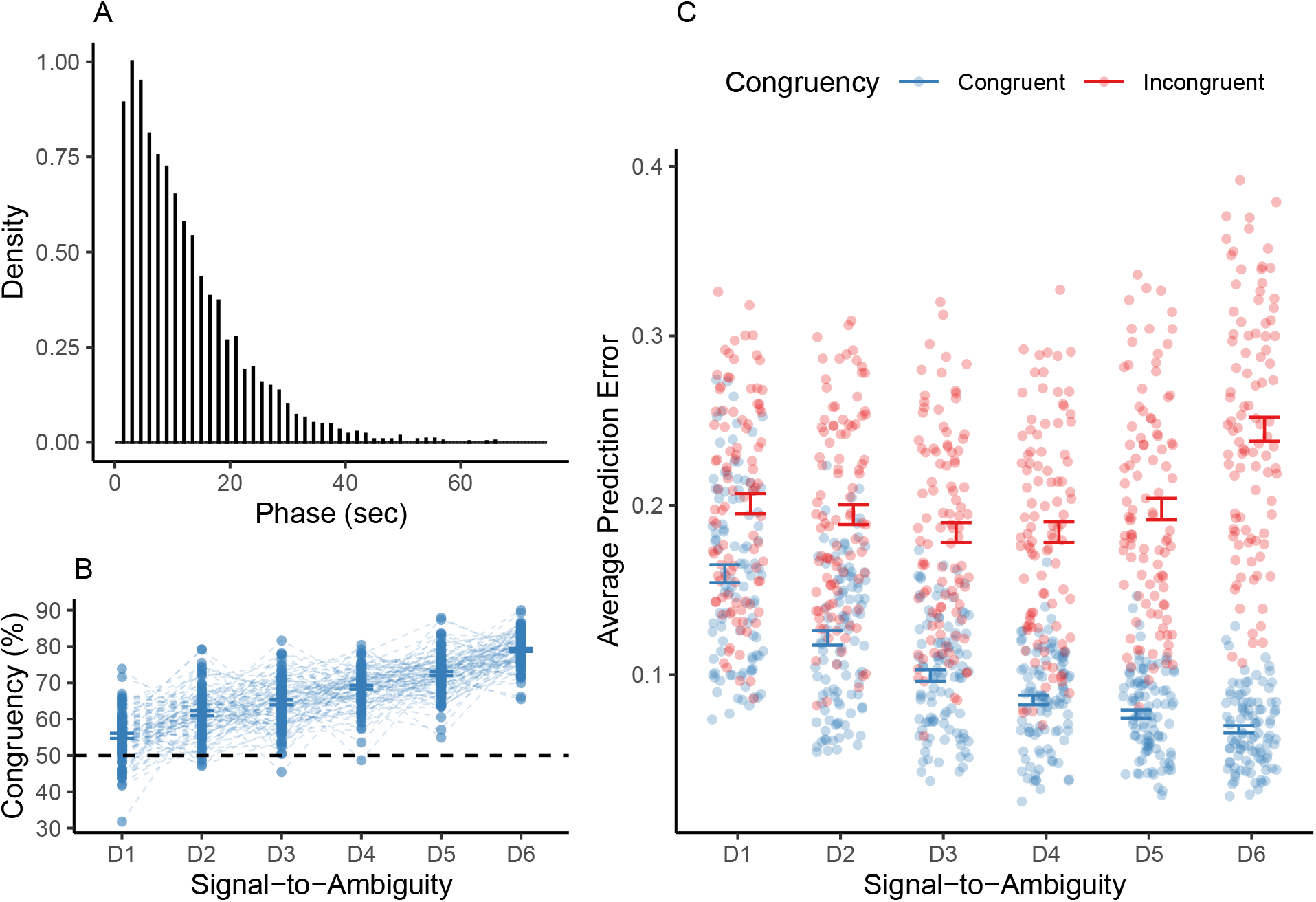
Simulating from the predictive-coding model of bistable perception. **A. Phase duration.** For ambiguous visual stimulation in 100 simulated participants, phase durations were distributed in a sharp rise and long tail. This pattern, which is typical for bistable perception, was best fit by a gamma distribution with a shape parameter of 1.66 and a rate parameter of 0.14 (BIC = 3.83 × 10^4^). **B. Graded ambiguity.** During graded ambiguity, disambiguating stimulus information biased simulated conscious experience toward congruent perceptual states. In analogy to our behavioral results, this simulated congruency effect was stronger at higher levels of signal-to-ambiguity (F(495) = 195.1, p = 1.49 × 10^-114^, *BF*_10_ = 4.18 × 10^120^). **C. Average prediction errors.** When simulating graded ambiguity, we found that prediction errors were elevated during incongruent as opposed to congruent perceptual states (main effect of *Congruency*: F(1.09 × 10^3^) = 4.15 × 10^3^, p = 0, *BF*_10_ = 7.16 × 10^275^). In analogy to our behavioral results, the modulation of simulated perceptual conflict was enhanced at higher signal-to-ambiguity ratios (interaction between *Congruency* and *Signal-to-Ambiguity*: F(1.09 × 10^3^) = 148.71, p = 2.1 × 10^-120^, *BF*_10_ = 1.99 × 10^116^). In contrast to our behavioral results, we also observed a main effect of *Signal-to-Ambiguity* (F(1.09 × 10^3^) = 81.43, p = 1.07 × 10^-72^, *BF*_10_ = 1.28 × 10^42^).

**Supplementary Figure S11.**
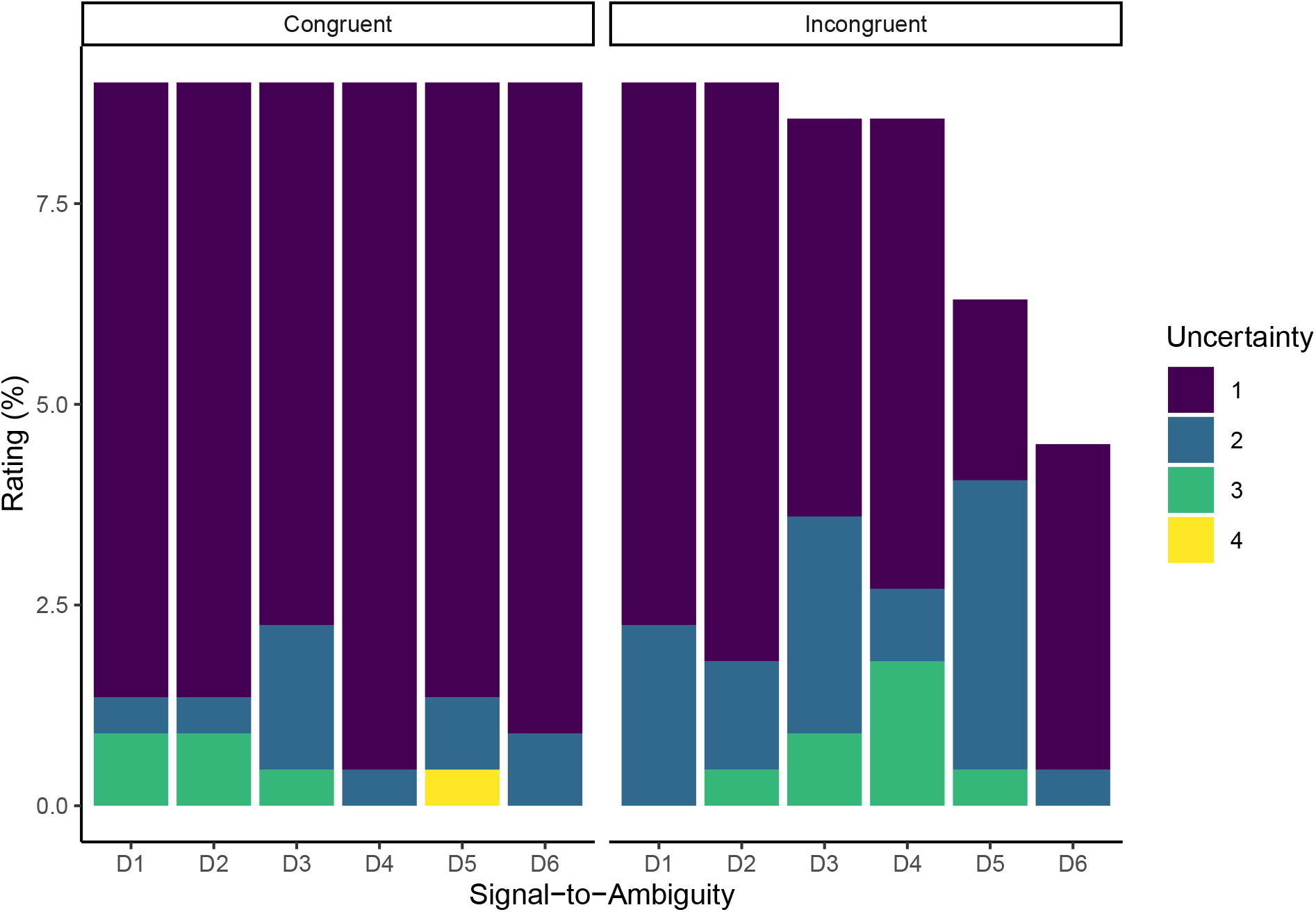
Perceptual uncertainty in E2. In the Run 3 of E2, participants viewed one run of graded ambiguity (six levels of signal-to-ambiguity D1-D6, each in one block of 120 sec visual stimulation) and indicated whether they perceived left-ward, rightward and unclear direction of rotation of the illusory sphere. In each block, we interrupted visual stimulation at random timepoints during both congruent and incongruent perceptual states. Participants rated perceptual uncertainty on a four-point scale (1: completely certain, 2: rather certain, 3: rather uncertain, 4: completely uncertain). The y-axis displays the percentage of uncertainty-reports recorded at every level of disambiguating stimulus information for both congruent and incongruent perceptual states. Uncertainty ratings were low during both congruency (average rating: 1.2 ± 0.05) and incongruency (1.4 ± 0.06) to disambiguating stimulus information. Ordinal repeated-measures ANOVA indicated a main effect of *Congruency* (Chi-square = 19.43, p = 3.5 × 10^-3^, *BF*_10_ = 27.3). Uncertainty ratings did not vary significantly with *Signal-to-Ambiguity* (Chi-square = 18.19, p = 0.05, *BF*_10_ = 2.99 × 10^-4^). Importantly, we also did not find an interaction between *Congruency* and *Signal-to-Ambiguity* (Chi-square = 7.41, p = 0.19, *BF*_10_ = 5.53 × 10^-5^). This argued against differences in perceptual uncertainty as a potential confound in our analysis of the neural correlates of graded ambiguity.

**Supplementary Figure S12.**
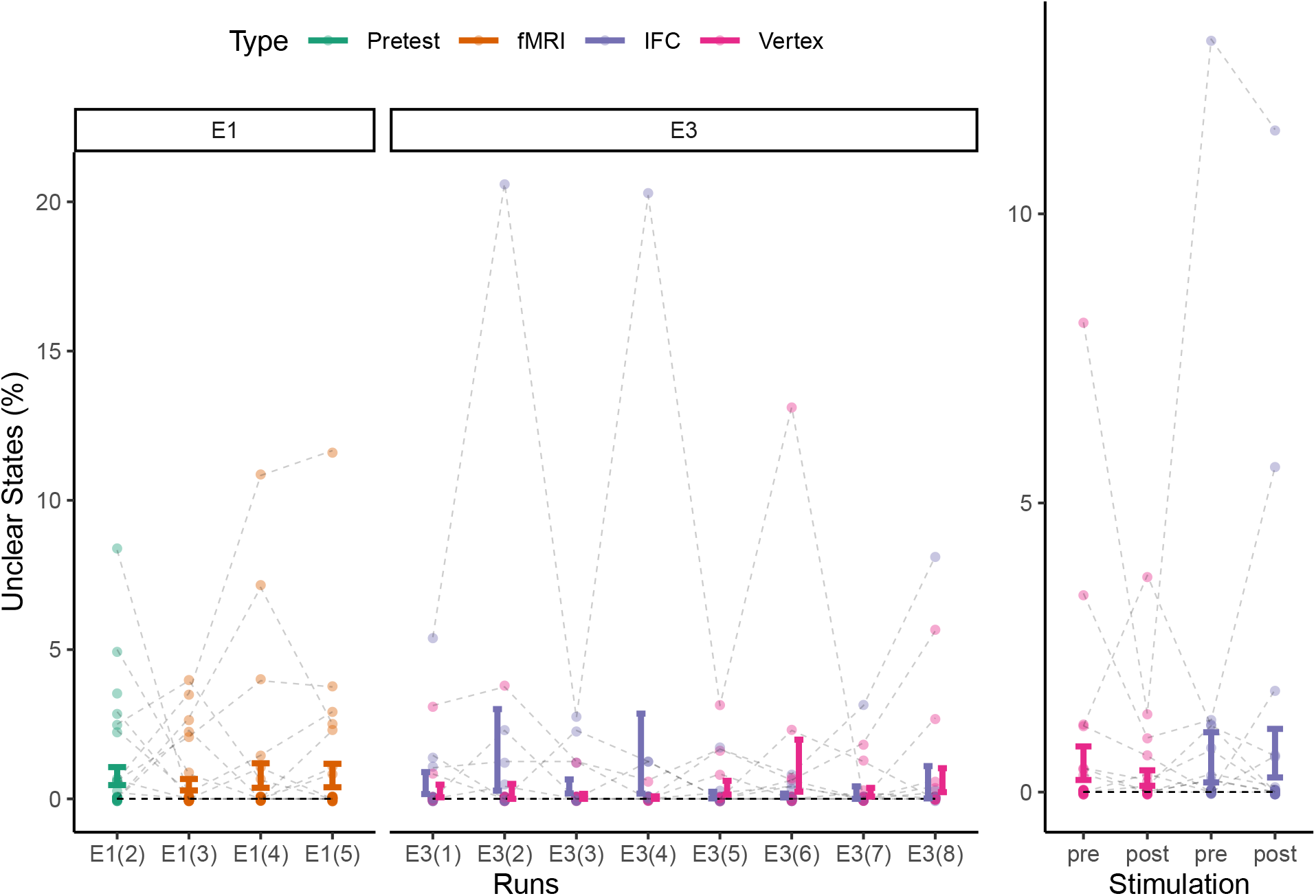
Unclear Perceptual States. Across E1 and E3, the frequency of unclear perceptual states did not vary with prolongated stimulus exposure (F(328.79) = 0.72, p = 0.72; left panel). In E3, we found no main effect of *Stimulation* (post vs. pre; F(58) = 0.35, p = 0.56, *BF*_10_ = 0.21), no main effect of *Session* (IFC vs. vertex; F(58) = 0.34, p = 0.56, *BF*_10_ = 0.41) and no between-factor interaction (F(58) = 1.12, p = 0.3, *BF*_10_ = 0.35; right panel).

### 9.10 Supplementary Tables

**Supplementary Table 1.**
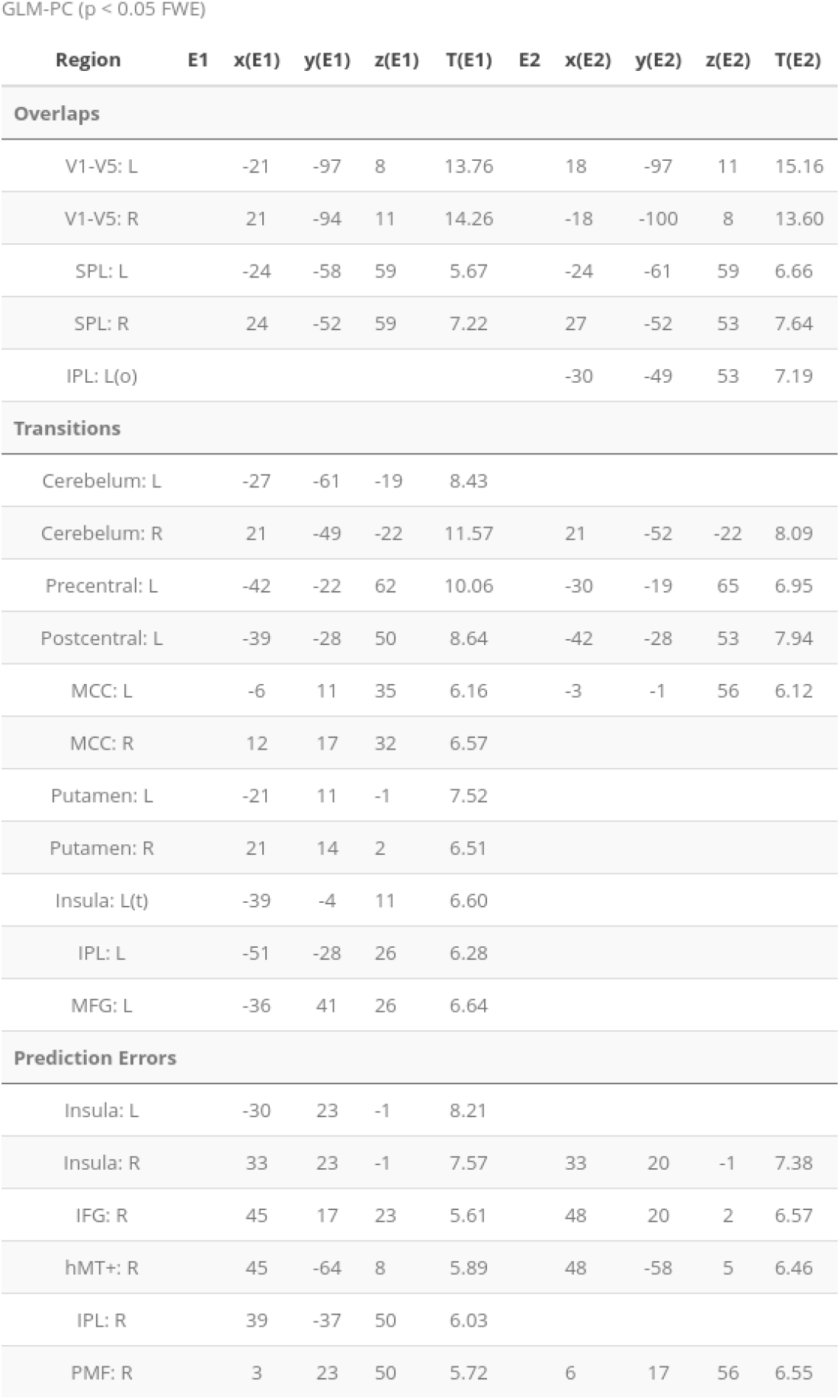
The neural correlates of perceptual conflict. Here, we summarize the fMRI results for GLM-PC (experiments E1 and E2, all *p_FWE_* < 0.05). Visual stimulation (“Overlaps”) correlated with BOLD signals throughout visual cortex (including V5/hMT+), bilateral superior parietal lobulus (SPL) and inferior parietal lobulus (IPS). At the time of perceptual events, we found *change-related* activations in bilateral cerebellum, left pre- and postcentral gyrus, bilateral midcingulate cortex and putamen, left insula, left IPL as well as left medial frontal gyrus (MFG). Perceptual prediction errors (i.e., the quantitative representation of perceptual conflict) correlated with BOLD signals in bilateral insula, right inferior frontal gyrus (IFG), right V5/hMT+, right IPL and right posterior-medial frontal gyrus (PMF; see corresponding maps in Figure 1C).

**Supplementary Table 2.** The neural correlates of graded ambiguity. To test the relation of V5/hMT+ and IFC to perceptual conflict during graded ambiguity, we modeled congruent and incongruent perceptual states by box-car regressors, separately for all six levels of signal-to-ambiguity. Group results were assessed using second-level ANOVA for the factors *Congruency* and *Signal-to-Ambiguity*. We found a main effect of *Congruency* (i.e., enhanced BOLD signals during incongruent as opposed to congruent perceptual states; *p_FWE_* < 0.05; see corresponding maps in Figure 3C) in right-hemispherical IFC (insula and inferior frontal gyrus), V5/hMT+, left precentral gyrus, right posterior-medial frontal gyrus (PMF) and right fusiform gyurs. We did not find any whole-brain correctable main effect of *Signal-to-Ambiguity*. Within the main effect of *Congruency* (thresholded at *p_FWE_* < 0.05), the interaction between *Congruency* and *Signal-to-Ambiguity* was significant in right-hemispherical IFC (insula and inferior frontal gyrus), V5/hMT+, left precentral gyrus, right PMF and right fusiform gyrus. Two additional cluster for this interaction in bilateral V1 (*p_FWE_* < 0.05) did not show a main effect of *Congruency*.

| Region | Congruency | x(M) | y(M) | z(M) | F(M) | Congruency x Signal-to-Ambiguity | x(I) | y(I) | z(I) | F(I) |
| --- | --- | --- | --- | --- | --- | --- | --- | --- | --- | --- |
| IFG: R |  | 45 | 23 | 29 | 31.30 |  | 45 | 17 | 26 | 5.69* |
| Insula: R |  | 33 | 23 | 5 | 23.14 |  | 33 | 23 | 2 | 5.30* |
| hMT+: R |  | 45 | -61 | 5 | 23.57 |  | 48 | -64 | 8 | 5.63* |
| Precentral: L |  | -42 | -4 | 38 | 35.05 |  | -42 | -4 | 35 | 5.54* |
| PMF: R |  | 6 | 17 | 50 | 31.66 |  | 6 | 14 | 47 | 5.87* |
| Fusiform: R |  | 33 | -67 | -10 | 24.20 |  | 33 | -67 | -10 | 4.67* |
| V1: L |  |  |  |  |  |  | -15 | -73 | 5 | 7.19 |
| V1: R |  |  |  |  |  |  | 15 | -70 | 8 | 8.23 |

## Notes

### Competing Interest Statement

The authors have declared no competing interest.

### Summary of Updates

This manuscript was updated following peer-review. We have revised the abstract, introduction, results and discussion section. In addition, we have added two new analyses (even-related BOLD timecourses and dynamic causal modeling) to the supplement.

## References

1. Michel, M. et al. Opportunities and challenges for a maturing science of consciousness. Nature Human Behaviour 3, 104–107 (2019).

2. Sohn, E. Decoding the neuroscience of consciousness. Nature 571, S2–S5 (2019).

3. Dehaene, S. et al. What is consciousness, and could machines have it? Science 358, 486–492 (2017).

4. Odegaard, B. et al. Should a Few Null Findings Falsify Prefrontal Theories of Conscious Perception? The Journal of Neuroscience 37, 9593–9602 (2017).

5. Boly, M. et al. Are the Neural Correlates of Consciousness in the Front or in the Back of the Cerebral Cortex? Clinical and Neuroimaging Evidence. The Journal of Neuroscience 37, 9603–9613 (2017).

6. Aru, J. et al. Distilling the neural correlates of consciousness. Neuroscience and Biobehavioral Reviews 36, 737–746 (2012).

7. Tsuchiya, N. et al. No-Report Paradigms: Extracting the True Neural Correlates of Consciousness. Tics 19, 757–770 (2015).

8. Brascamp, J. et al. Multistable Perception and the Role of the Frontoparietal Cortex in Perceptual Inference. Annual Review of Psychology 69, 77–103 (2018).

9. Blake, R. et al. Visual competition. Nature Reviews Neuroscience 3, 13–21 (2002).

10. Leopold, D. A. et al. Multistable phenomena: changing views in perception. Tics 3, 254–264 (1999).

11. Hohwy, J. et al. Predictive coding explains binocular rivalry: an epistemological review. Cognition 108, 687–701 (2008).

12. Sterzer, P. et al. The neural bases of multistable perception. Tics 13, 310–8 (2009).

13. Lumer, E. D. et al. Neural correlates of perceptual rivalry in the human brain. Science 280, 1930–4 (1998).

14. Weilnhammer, V. et al. A predictive coding account of bistable perception - a model-based fMRI study. PLOS Computational Biology 13, e1005536 (2017).

15. Knapen, T. et al. The Role of Frontal and Parietal Brain Areas in Bistable Perception. The Journal of neuroscience 31, 10293–10301 (2011).

16. Frässle, S. et al. Binocular rivalry: frontal activity relates to introspection and action but not to perception. The Journal of neuroscience 34, 1738–47 (2014).

17. Brascamp, J. et al. Negligible fronto-parietal BOLD activity accompanying unreportable switches in bistable perception. Nature neuroscience 18, 1672–1678 (2015).

18. Zou, J. et al. Binocular rivalry from invisible patterns. PNAS 113, 8408–8413 (2016).

19. Pastukhov, A. et al. Believable change: bistable reversals are governed by physical plausibility. Journal of vision 12, (2012).

20. Knill, D. C. et al. The Bayesian brain: the role of uncertainty in neural coding and computation. Trends Neurosci. 27, 712–719 (2004).

21. Hohwy, J. Attention and conscious perception in the hypothesis testing brain. Frontiers in psychology 3, 96 (2012).

22. Rosa, M. J. et al. Bayesian model selection maps for group studies. NeuroImage 49, 217–24 (2010).

23. Heekeren, H. R. et al. A general mechanism for perceptual decision-making in the human brain. Nature 431, 859–862 (2004).

24. Krug, K. et al. A causal role for V5/MT neurons coding motion-disparity conjunctions in resolving perceptual ambiguity. Current Biology 23, 1454–1459 (2013).

25. Friston, K. J. et al. Dynamic causal modelling. NeuroImage 19, 1273–1302 (2003).

26. Haynes, J. D. et al. Decoding mental states from brain activity in humans. Nature Reviews Neuroscience 7, 523–534 (2006).

27. Kersten, D. et al. Object Perception as Bayesian Inference. Annual Review of Psychology 55, 271–304 (2004).

28. Weilnhammer, V. et al. Psychotic Experiences in Schizophrenia and Sensitivity to Sensory Evidence. Schizophrenia bulletin (2020).

29. Van Gaal, S. et al. Unconscious activation of the prefrontal no-go network. Journal of Neuroscience 30, 4143–4150 (2010).

30. Huang, Y. Z. et al. Theta burst stimulation of the human motor cortex. Neuron 45, 201–206 (2005).

31. Kriegeskorte, N. et al. Information-based functional brain mapping. PNAS 103, 3863–3868 (2006).

32. Schmack, K. et al. Predicting Subjective Affective Salience from Cortical Responses to Invisible Object Stimuli. Cerebral cortex 26, 3453–3460 (2016).

33. Wilson, H. R. Minimal physiological conditions for binocular rivalry and rivalry memory. Vision research 47, 2741–50 (2007).

34. Xu, H. et al. Rivalry-like neural activity in primary visual cortex in anesthetized monkeys. The Journal of Neuroscience 36, 3231–3242 (2016).

35. Corbetta, M. et al. The Reorienting System of the Human Brain: From Environment to Theory of Mind. vol. 58 306–324 (2008).

36. Baldauf, D. et al. Neural mechanisms of object-based attention. Science 344, 424–427 (2014).

37. Aron, A. R. et al. Inhibition and the right inferior frontal cortex: One decade on. Tics 18, 177–185 (2014).

38. Alais, D. et al. Attending to auditory signals slows visual alternations in binocular rivalry. Vision Research 50, 929–935 (2010).

39. Prado, J. et al. Variations of response time in a selective attention task are linked to variations of functional connectivity in the attentional network. NeuroImage 54, 541–549 (2011).

40. Dwarakanath, A. et al. Prefrontal state fluctuations control access to consciousness. bioRxiv 2020.01.29.924928 (2020) doi:10.1101/2020.01.29.924928.

41. Dürschmid, S. et al. Direct Evidence for Prediction Signals in Frontal Cortex Independent of Prediction Error. Cerebral Cortex 29, 4530–4538 (2019).

42. Klink, P. C. et al. Early interactions between neuronal adaptation and voluntary control determine perceptual choices in bistable vision. Journal of Vision 8, 16 (2008).

43. Graaf, T. A. de et al. On the functional relevance of frontal cortex for passive and voluntarily controlled bistable vision. Cerebral cortex (New York, N.Y.: 1991) 21, 2322–31 (2011).

44. Stephan, K. E. et al. Nonlinear dynamic causal models for fMRI. NeuroImage 42, 649–662 (2008).

45. Toppino, T. C. et al. Time for a change: What dominance durations reveal about adaptation effects in the perception of a bi-stable reversible figure. Attention, Perception, and Psychophysics 77, 867–882 (2015).

46. Moreno-Bote, R. et al. Noise-Induced Alternations in an Attractor Network Model of Perceptual Bistability. Journal of Neurophysiology 98, 1125–1139 (2007).

47. Garrido, M. I. et al. The mismatch negativity: A review of underlying mechanisms. Clinical Neurophysiology 120, 453–463 (2009).

48. Corlett, P. R. et al. Hallucinations and Strong Priors. Trends in cognitive sciences 23, 114–127 (2019).

49. Sommer, I. E. C. et al. Auditory verbal hallucinations predominantly activate the right inferior frontal area. Brain 131, 3169–77 (2008).

50. Powers, A. R. et al. Pavlovian conditioning–induced hallucinations result from over-weighting of perceptual priors. Science 357, 596–600 (2017).

51. Wang, M. et al. Brain mechanisms for simple perception and bistable perception. Proceedings of the National Academy of Sciences of the United States of America 110, E3350–E3359 (2013).

52. Panagiotaropoulos, T. I. et al. Neuronal Discharges and Gamma Oscillations Explicitly Reflect Visual Consciousness in the Lateral Prefrontal Cortex. Neuron 74, 924–935 (2012).

53. Kapoor, V. et al. Decoding the contents of consciousness from prefrontal ensembles. bioRxiv 2020.01.28.921841 (2020) doi:10.1101/2020.01.28.921841.

54. Hesse, J. K. et al. A new no-report paradigm reveals that face cells encode both consciously perceived and suppressed stimuli. eLife 9, (2020).

55. Brainard, D. H. The Psychophysics Toolbox. Spatial vision 10, 433–6 (1997).

56. Weilnhammer, V. A. et al. Frontoparietal cortex mediates perceptual transitions in bistable perception. The Journal of neuroscience 33, 16009–15 (2013).

57. Morgan, M. J. et al. Apparent motion and the Pulfrich effect. Perception 4, 3–18 (1975).

58. O’Shea, J. et al. Transcranial magnetic stimulation. Current Biology 17, R196–R199 (2007).

59. Eickhoff, S. B. et al. A new SPM toolbox for combining probabilistic cytoarchitectonic maps and functional imaging data. NeuroImage 25, 1325–1335 (2005).

60. Rossini, P. M. et al. Non-invasive electrical and magnetic stimulation of the brain, spinal cord, roots and peripheral nerves: Basic principles and procedures for routine clinical and research application: An updated report from an I.F.C.N. Committee. Clinical Neurophysiology 126, 1071–1107 (2015).

61. Schicktanz, N. et al. Continuous theta burst stimulation over the left dorsolateral prefrontal cortex decreases medium load working memory performance in healthy humans. PloS one 10, e0120640 (2015).

62. Suppa, A. et al. Theta burst stimulation induces after-effects on contralateral primary motor cortex excitability in humans. Journal of Physiology 586, 4489–4500 (2008).

63. Valchev, N. et al. Primary somatosensory cortex necessary for the perception of weight from other people’s action: A continuous theta-burst TMS experiment. NeuroImage 152, 195–206 (2017).

64. Weilnhammer, V. A. et al. Revisiting the Lissajous figure as a tool to study bistable perception. Vision research 98, 107–12 (2014).

65. Friston, K. A theory of cortical responses. Philosophical transactions of the Royal Society of London. Series B, Biological sciences 360, 815–836 (2005).

66. Friston, K. J. et al. Free-energy and the brain. Synthese 159, 417–458 (2007).

67. Hebart, M. N. et al. The Decoding Toolbox (TDT): a versatile software package for multivariate analyses of functional imaging data. Frontiers in Neuroinformatics 8, 88 (2015).

68. Cortes, C. et al. Support-vector networks. Machine Learning 20, 273–297 (1995).

69. Suzuki, S. et al. Long-Term Speeding in Perceptual Switches Mediated by Attention-Dependent Plasticity in Cortical Visual Processing. Neuron 56, 741–753 (2007).

70. Ostermann, T. et al. Regression toward the mean – a detection method for unknown population mean based on Mee and Chua’s algorithm. BMC Medical Research Methodology 8, 52 (2008).

71. Stephan, K. E. et al. Bayesian model selection for group studies. NeuroImage 46, 1004–17 (2009).

